# Combinatorial patterns of gene expression changes contribute to variable expressivity of the developmental delay-associated 16p12.1 deletion

**DOI:** 10.1101/2021.03.06.434203

**Authors:** Matthew Jensen, Anastasia Tyryshkina, Lucilla Pizzo, Corrine Smolen, Maitreya Das, Emily Huber, Arjun Krishnan, Santhosh Girirajan

## Abstract

**Background:** Recent studies have suggested that individual variants do not sufficiently explain the variable expressivity of phenotypes observed in complex disorders. For example, the 16p12.1 deletion is associated with developmental delay and neuropsychiatric features in affected individuals, but is inherited in >90% of cases from a mildly-affected parent. While children with the deletion are more likely to carry additional “second-hit” variants than their parents, the mechanisms for how these variants contribute to phenotypic variability are unknown.

**Methods:** We performed detailed clinical assessments, whole-genome sequencing, and RNA sequencing of lymphoblastoid cell lines for 32 individuals in five large families with multiple members carrying the 16p12.1 deletion. We identified contributions of the 16p12.1 deletion and “second-hit” variants towards a range of expression changes in deletion carriers and their family members, including differential expression, outlier expression, alternative splicing, allele-specific expression, and expression-quantitative trait loci analyses.

**Results:** We found that the deletion dysregulates multiple autism and brain development genes such as *FOXP1*, *ANK3*, and *MEF2*. Carrier children also showed an average of 5,323 gene expression changes compared with one or both parents, which matched with 33/39 observed developmental phenotypes. We identified significant enrichments for 13/25 classes of “second-hit” variants in genes with expression changes, where 4/25 variant classes were only enriched when inherited from the non-carrier parent, including loss-of-function SNVs and large duplications. In 11 instances, including for *ZEB2* and *SYNJ1*, gene expression was synergistically altered by both the deletion and inherited “second-hits” in carrier children. Finally, brain-specific interaction network analysis showed strong connectivity between genes carrying “second-hits” and genes with transcriptome alterations in deletion carriers.

**Conclusions:** Our results suggest a potential mechanism for how “second-hit” variants modulate expressivity of complex disorders such as the 16p12.1 deletion through transcriptomic perturbation of gene networks important for early development. Our work further shows that family-based assessments of transcriptome data are highly relevant towards understanding the genetic mechanisms associated with complex disorders.

## BACKGROUND

Complex disorders, such as autism, intellectual disability/developmental delay (ID/DD), epilepsy, and schizophrenia, have been attributed to rare copy-number variants (CNVs), or deletions and duplications encompassing multiple genes, as well as individual rare single nucleotide variants (SNVs) and the combined effects of common variants (1–6). Despite advances in high-throughput sequencing methods and quantitative assessments of large cohorts, individual variants implicated for these disorders do not sufficiently explain the variable expressivity and pleiotropy of clinical features often observed in affected individuals (7–9). An example is the 520-kbp deletion at chromosome 16p12.1 (OMIM: 136570), which was originally described in children with developmental delay (10, 11) but was subsequently found to confer increased risk for schizophrenia (12), epilepsy (13), and cognitive defects in control populations (14). Unlike syndromic CNVs such as Smith-Magenis syndrome that primarily occur *de novo* (15), the 16p12.1 deletion is inherited in over 90% of affected children from carrier parents who manifest subclinical or mild cognitive and neuropsychiatric features (10, 16). In fact, we recently found that children with the deletion were more likely to carry an additional burden of rare CNVs (11) and deleterious variants in genes intolerant to variation (16), or “second-hit” variants, compared to their carrier parents, making the deletion an ideal model for assessing the combined effects of multiple variants towards variable clinical outcomes.

While dissecting the pathogenicity of complex disorders has been challenging, cohort and family-based studies that integrate multiple variants with different effect sizes or functional outcomes have provided insights into how the genetic architecture contributes to changes in penetrance, severity, and complexity of phenotypes. In particular, analysis of gene expression patterns in human cells allows for dissecting the direct and indirect effects of genomic variants towards biological functions in complex disorders. *For example*, Merla and colleagues assessed gene expression in skin fibroblasts and lymphoblastoid cell lines (LCLs) from individuals with the 7q11.23 deletion, associated with Williams syndrome, and found that several genes adjacent to the deletion region were also downregulated compared to controls (17). Similarly, expression changes due to the autism-associated 16p11.2 deletion correlated with changes in head circumference phenotypes (18) and converged on several neurodevelopmental pathways, including synaptic function and chromatin modification (19). Other studies have used expression data to identify pathogenic variants potentially missed by genome sequencing studies (20–22). *For example*, Frésard and colleagues identified novel causal variants for 6/80 individuals with rare undiagnosed diseases through paired analysis of whole-blood transcriptomes and genomes (20). Additionally, several recent studies have used family-based approaches to study the effects of rare inherited variants towards gene expression. *For example,* Pala and colleagues found that rare inherited variants in both coding and non-coding regions increased the likelihood of gene expression changes among 61 families in the bottlenecked Sardinia population, indicating the importance of such variants towards disease risk (23). While these studies have shown the utility of assessing transcriptomic consequences of individual causal variants, they were focused on either control populations *or* relatively invariable disorders, and did not examine the simultaneous effects of multiple variants with different effect sizes towards changes in gene expression within major biological pathways.

Here, we integrated whole-genome sequencing and transcriptome data of LCLs from 32 individuals in five large 16p12.1 deletion families who manifested variable phenotypes, in order to investigate how the combined effects of the deletion and “second-hits” perturb transcriptional networks and biological functions. We found that the 16p12.1 deletion disrupts expression of genes involved in neuronal and developmental functions, such as signal transduction and cell proliferation, as well as genes preferentially expressed in the fetal and adult brain. We further identified significant contributions of several classes of rare “second-hit” coding and non-coding variants towards changes in gene expression among carrier children compared with their parents, especially when the variants were inherited from the noncarrier parent. In fact, we found 11 instances of genes in carrier children whose expression was synergistically altered by the combined effects of the 16p12.1 deletion and “second-hit” variants inherited from the noncarrier parent. Although a relatively small sample size precluded global analyses between these expression changes and developmental phenotypes, we found that specific expression changes contributed towards distinct clinical features of affected children through disruption of biological functions related to neurodevelopment. Our results suggest that the 16p12.1 deletion and “second-hit” variants jointly disrupt the developmental transcriptome through shared pathways to contribute towards developmental phenotypes, emphasizing the importance of family-based transcriptome studies for complex disorders.

## METHODS

### Patient recruitment and clinical phenotype analysis

We obtained clinical data and whole-blood DNA from 32 individuals in five families with the 16p12.1 deletion. Among the recruited individuals were 10 children with the deletion (“carrier children”), six sets of carrier and noncarrier parents (including one family with two pairs of parents), three sets of carrier and noncarrier grandparents, and four noncarrier siblings (**Figure S1**; **Table S1**). Affected children and family members were identified as carriers of the deletion through prior clinical diagnostic tests, which we confirmed using SNP microarray analysis (16). We collected phenotypic information from the five families using two standardized clinical questionnaires: one assessing developmental phenotypes in children, and the other assessing psychiatric features in adults. These data represent comprehensive medical history of affected children and their family members, including neuropsychiatric and developmental features (including cognitive, behavioral, and psychiatric diagnoses), anthropomorphic measures, abnormalities across multiple organ systems (nervous, craniofacial, musculoskeletal, cardiac, hearing/vision, digestive, and urinary systems), and family history of medical or psychiatric disorders. Family members first submitted completed checklists eliciting major phenotypes and medical history, which were then integrated with detailed medical records for each person. A follow-up phone interview was then conducted with family members to fill in any missing information on the clinical questionnaire. Using this method, we assessed clinical data on 31/32 individuals in the cohort. Summarized clinical features for children and adults in this study are listed in **Table S1**. We note that all families had self-reported European or Caucasian ancestry. Based on the curated phenotypic data, we calculated quantitative scores for children using a modified de Vries scoring rubric, as described previously (16), which represents the diversity and severity of phenotypic features in affected children (24). We similarly summed the number of neuropsychiatric features to generate phenotypic scores in adults. Phenotypic scores for all individuals in the cohort are listed in **Table S1**.

### DNA extraction, whole-genome sequencing, and variant identification

We identified 25 classes of rare deleterious variants from whole-genome sequencing (WGS) and SNP microarray for each of the 32 family members in our cohort. The 25 rare variant classes identified in this study are displayed in **Figure S2** and listed in **Table S2**. Genomic DNA was extracted from peripheral blood using QIAamp DNA Blood Maxi extraction kit (Qiagen, Hilden, Germany) and treated with RNAse. DNA levels were then quantified using Quant-iT™ PicoGreen™ dsDNA assay methods (Thermo Fisher Scientific, Waltham, MA, USA), and sample integrity was assessed in agarose gel. After constructing Illumina TruSeq DNA PCR-free libraries (San Diego, CA, USA), whole-genome sequencing was performed on each sample by Macrogen Labs (Rockville, MD, USA) using an Illumina HiSeq X sequencer to obtain an average coverage of 34.5X. Raw sequencing data were processed for quality control using Trimmomatic (25) with leading:5, trailing:5, and slidingwindow:4:20 parameters, aligned to the human hg19 reference genome using BWAv.0.7.13 (26), and sorted and indexed using Samtools v.1.9 (27).

The GATK Best Practices pipeline v.3.8 (HaplotypeCaller) and v.4.0.11 (GenotypeGVCFs) (28) was used to identify SNVs and small indels from WGS data. In short, duplicate reads were marked and removed using PicardTools, and after calibration of base-pair quality scores, GATK HaplotypeCaller was used to identify variants in each sample. Variant calls were then pooled for joint genotyping and calibration of variant quality scores. Custom-built pipelines using Annovar v.2016Feb01 (29) applied a total of 430 annotation classes for variant function, population frequency, conservation, genomic region, and predicted pathogenicity. Variants were filtered based on the following quality metrics (30): QUAL >50, read depth >8, allele balance between 0.25-0.75 (or >0.9 for homozygous variants), and quality depth (QUAL/reads with alternate allele) <1.5. Rare variants were defined as variants with frequency ≤0.001 in the gnomAD v.2.1.1 genome database (31), and present in <10 samples in our in-house WGS cohort of 125 families (335 individuals) with rare CNVs, in order to remove technical artifacts that may be missed by gnomAD. We finally classified rare SNVs and small indels for downstream analysis as follows: Rare missense and loss-of-function (LOF, including frameshift and stopgain) variants within protein coding regions, as well as variants in the 5’ and 3’ untranslated region (UTR) or within 1 kbp of the transcription start (TSS) or end sites (upstream and downstream), were classified based on their RefSeq-defined genomic locations in Annovar (**Figure S2**). Splice-site variants were identified based on MutationTaster annotations (32) for disease-causing (“D”) or disease-causing automatic (“A”) variants. Rare non-coding regulatory variants within 50 kbp of TSS for protein-coding genes were classified according to ChromHMM chromatin state segmentation data for GM12878 lymphoblastoid cells (33), available from the ENCODE Project, into promoters (chromosome states 1-3), enhancers (states 4-7), or silencers (state 12). With the exception of loss-of-function and splice-site variants, all coding and non-coding variants were filtered for CADD Phred-like pathogenicity scores ≥10 (34). Inheritance patterns of these variants were determined using in-house pipelines.

Copy-number variants and structural variants were identified using a combination of WGS data for all samples and SNP microarray data for 25/32 samples, previously described in (16). To identify variants from microarray data, Illumina Omni 2.5 BeadChip microarray experiments were performed for each sample at either the HudsonAlpha Institute for Biotechnology (Huntsville, AL, USA; n=18), Yale Center for Genome Analysis (New Haven, CT, USA; n=5), or the Department of Genome Sciences at the University of Washington (Seattle, WA, USA; n=2). CNV calls for each sample were generated using PennCNV v.1.0.3 (35), and were filtered for ≥50 kbp in length and ≥5 target probes. CNVs and SVs were also detected from aligned WGS data using a combination of four pipelines: CNVNator v.0.4.1 (36) (bin size of 200), DELLYv.0.8.2 (37), LUMPY-sv v.0.2.13 with Smoove v.0.2.5 (38), and Manta v.1.6.0 (39). In both WGS and microarray-derived datasets, adjacent CNVs were merged if they overlapped or had a gap <20% of CNV length and <50 kbp. We then integrated the CNV and SV calls from each of the datasets as follows: For smaller CNVs and SVs <50 kbp, any duplication or deletion called by at least two of the four WGS-based callers were considered for downstream analysis, with the minimum intersected regions defining the new breakpoints. For larger CNVs and SVs >50 kbp, the union of CNVNator and PennCNV calls were considered for downstream analysis. Integrated calls were based on 50% reciprocal overlap among the callers. As our SV call-set had a low overlap with SV call-sets from control populations, likely due to different SV calling methods used in the control cohorts (40, 41), integrated variants were filtered for presence in <10 individuals in our in-house WGS cohort, as determined by 50% reciprocal overlap. Finally, RefSeq gene-coding regions spanned by SVs were categorized as follows: encapsulating variants which span the entire gene, interstitial variants that are contained within a gene, and 5’ and 3’ UTR variants that overhang the gene on either end (**Figure S2**). Inheritance patterns of CNV and SV calls were determined if calls in the child and parent had >50% reciprocal overlap.

Short tandem repeats (STRs) were called from aligned WGS data with GangSTR v.2.4 (42), using the GangSTR hg19 reference file v.13.1. The calls were filtered and analyzed using three tools from the STR analysis toolkit TRTools (43). First, dumpSTR was used to filter for quality of calls using the following parameters: read depth >20, read depth <1000, QUAL >0.9, spanbound only (calls that are spanned by reads), and filter bad confidence intervals (filtered calls whose maximum likelihood estimates were not within the confidence interval). The reads were then merged with mergeSTR, and basic statistics were calculated using statSTR. In addition to the dumpSTR filters, we applied the following filters to our call set: >95% of samples called for the STR location, variance at location >0, and overlap of the STR location with a RefSeq defined protein-coding gene. STR expansions were defined as any call for which the deviation of the repeat length was greater than the mean length plus three standard deviations among all individuals in our cohort. Finally, STR variants were categorized according to their genomic location, including exonic, intronic, 5’ or 3’ UTR, upstream, and downstream (**Figure S2**). Inheritance patterns of STRs were determined by matching the number of repeats in the child to their parents.

### Generation of lymphoblastoid cell lines and RNA-sequencing

Peripheral blood samples for all 32 individuals in our cohort were submitted to the Coriell Institute for Medical Research (Camden, NJ, USA) for generation of lymphoblastoid cell lines through Epstein-Barr virus transformation of B-lymphocytes (**Table S3**). After receiving the LCL samples, cells were grown at 5% CO2 and a concentration of 1X 106 cells/mL under L-glutamine-supplemented RPMI 1640 medium (11875-119, Thermo Fisher Scientific) containing 15% fetal bovine serum (35-010-CV, Corning Life Sciences, Tewksbury, MA, USA) and Cytiva HyClone™ Penicillin Streptomycin solution (SV30010, Thermo Fisher Scientific). Total RNA was isolated from three biological replicates of P6-P7 cells per sample using TRIzol Reagent (Thermo Fisher Scientific) and PureLink RNA Mini Kit (12183018A, Thermo Fisher Scientific), and subsequently treated with DNA-free DNA Removal Kit (AM1906, Thermo Fisher Scientific). RNA integrity number scores (RIN) were assessed using Agilent Bioanalyzer 2100 (**Figure S3A**), and replicates with RIN scores >8.5 were sequenced. Paired-end 50 bp libraries for each replicate were generated using Illumina TruSeq Stranded mRNA kit, and were sequenced using Illumina NovaSeq at the Penn State College of Medicine Genome Sciences Facility (Hershey, PA, USA). Two sequencing runs of 48 replicates were performed, with the biological replicates of each sample split among the two runs to mitigate batch effects, to generate a total of 43.5 million reads/replicate.

### Quantification of gene expression and coverage of disease genes

Sequenced RNA reads were filtered using Trimmomatic v.0.36 (25) to remove reads <30 bp long. Following the GTEx Consortium RNA-Seq pipeline (44), the filtered reads were aligned to the human genome (GENCODE v.19) using STAR v.2.4.2a (45), and sorted and indexed using Samtools v.1.9 (27). Duplicates reads were marked with PicardTools v.2.9.0. We assessed the quality of the aligned reads with transcript integrity scores(46), which moderately correlated (r=0.38, p=1.0×10^-4^, Pearson correlation) with the RIN scores for each sample (**Figure S3A**). Isoform counts for GENCODE 19 genes were quantified using RSEM v.1.2.22 (47). A collapsed gene coordinate GTF file was generated using the GENCODE 19 gene coordinates and the GTEX collapse_annotation script. Gene-level counts and transcripts per million read (TPM) values were quantified using RNASeQC v.1.1.8 (48), using strict mode and the collapsed gene coordinates.

After filtering for transcripts where all three replicates of at least one sample showed >0.2 TPM, we obtained a total of 24,340 expressed transcripts across our cohort, representing 43.3% of all GENCODE transcripts. We further compared our set of expressed LCL genes to disease gene databases (49–53) and genes expressed in the adult brain from GTEx consortium RNA-Seq data (44). We defined expressed genes in GTEx tissues if they showed >0.5 TPM in 80% of samples for a particular tissue. The expressed LCL genes covered >70% of each of these gene sets, including 83% of genes expressed in GTEx brain tissues (**Figure S4A-B**). These data are in concordance with gene expression data from GTEx, where gene expression values in LCLs and brain tissues showed an average Spearman correlation of 0.84 (**Figure S4C**). These findings indicate that our LCL data would be able to identify changes in expression patterns for most genes related to neurodevelopmental disease.

### Differential expression and outlier expression analysis

We performed differential expression analysis between all 16p12.1 deletion carriers and noncarriers, as well as between parents and offspring across the five families, using edgeR(54) v.3.30.0 on gene-level counts to create generalized linear models and perform quasi-likelihood tests with Benjamini-Hochberg correction. For testing differences between all deletion carriers (n=19) and noncarriers (n=13), we included family as a covariate in the linear model, used default filtering for low-expressed genes, and removed genes with sex-specific differences in GTEx LCL samples as well as genes on the X and Y chromosomes (due to unequal sex ratios in deletion carriers and noncarriers). To control for expression outliers, we iteratively identified sets of differentially expressed genes, defined using an FDR<0.05 threshold (Benjamini-Hochberg correction), between deletion carriers and noncarriers after removing one sample at a time. We then took the intersection of the resulting 32 sets of differentially expressed genes, and obtained a total of 1,569 transcripts differentially expressed in the deletion carriers (**Table S4**). We also performed differential expression analysis using PQLseq v.1.2 to account for gene expression similarity due to relatedness (55). We first generated input files from unfiltered WGS SNV data using PLINK v.1.9 (56), and used GEMMA v.0.98.3, which calculates kinship between two individuals based on genotype similarity (57), to generate a kinship matrix for our cohort. This matrix was used as input for PQLseq along with gene-level counts from RNA-Seq data, after removing the same sex-specific genes as for the edgeR analysis.

We next performed family-based analysis on 13 separate trios identified across the five families (nine carrier children compared to parents and four carrier parents compared to grandparents), which are listed in **Table S1**. For example, we separately analyzed two trios in family GL_001 (**Figure S1**). For comparison, we analyzed an additional four trios with noncarrier children compared to carrier and noncarrier parents (**Figure S5C**). For each trio, we first used an edgeR workflow without covariates to identify differentially expressed genes between the offspring and carrier parent (|logFC|>0.5, FDR<0.05, Benjamini-Hochberg correction), and separately assessed expression changes between the offspring and noncarrier parent. Genes with low expression (expressed in <25% of all replicates) and sex-specific genes were removed from edgeR analysis. We then overlapped the two sets of differentially expressed genes to classify expression changes by family-specific patterns as follows: “unique” if the gene was differentially expressed in the offspring compared with both parents; “shared with the carrier parent” if the gene was only differentially expressed compared with the noncarrier parent; and “shared with the noncarrier parent” if the gene was only differentially expressed compared with the carrier parent (**Figure S5A**).

To identify genes with outlier expression in our cohort, we calculated z-scores of gene expression values for each individual for 14,212 protein coding genes expressed in the LCL samples. We normalized the expression values in each person by calculating the median TPM expression across the three replicates for each gene, transformed the values using log2(x + 1), and calculated z-scores for each log-transformed TPM compared with all samples in our cohort. As principal component analysis showed clustering of samples by family (**Figure S6A**), we used PEER v.1.0 (58) to correct the z-scores using one PEER principal component (**Figure S6B**). After correction, we further assessed for clustering of samples and replicates, and found strong Spearman correlations among replicates derived from the same sample (**Figure S3B**). We defined outlier genes as any gene with |z-score| >2 (**Figure S6C**), in line with recent studies utilizing outlier expression values (20).

### Enrichment analysis for biological function, brain expression, and disease relevance

Enrichment analysis for sets of differentially expressed genes was performed using goseq v.3.12 (59), which tests for overrepresentation of gene categories in RNA-seq data. Goseq controls for selection bias in RNA-seq datasets by modeling the distribution of transcript lengths of differentially expressed genes. We assessed for enriched biological processes using the Gene Ontology database (60), as well as genes expressed in specific adult brain tissues from GTEx (44) and developing brain tissues from the BrainSpan Atlas (61). We defined preferentially-expressed genes in GTEx and BrainSpan tissues as expression >2 standard deviations higher than the median expression across all tissues for that gene. We further assessed for enrichment of differentially expressed gene sets for candidate neurodevelopmental disease genes (DBD Gene Database) (49), as well as specific gene sets for autism (SFARI Gene database) (50), intellectual disability (DDD and DDG2P databases) (52, 53), and schizophrenia (51). Finally, we annotated sets of genes with altered expression for two common measures of intolerance to variation, RVIS (62) and pLI (63), and used genes considered to be intolerant to variation (RVIS <20^th^ percentile or pLI score >0.9) for downstream analysis. All gene sets used for enrichment analyses were filtered for genes with transcripts that are expressed in our LCL samples (>0.2 TPM in all three replicates of at least one sample).

### PAGE and WGCNA analysis in deletion carriers and noncarriers

We performed parametric analysis of gene set enrichment (PAGE) on genes that were differentially expressed between carriers and non-carriers of the 16p12.1 deletion (64). PAGE is a gene set analysis method that considers the direction of the expression log fold change to discover sets of genes that are enriched among up- or down-regulated genes. For this analysis, we included the log-fold change of 26,861 transcripts that were not filtered out by edgeR’s default filtering of low-expressed transcripts. We searched for significant up- or down-regulation of genes within terms from the Gene Ontology database (60), using two-tailed z-tests with Benjamini-Hochberg correction.

We further performed weighted gene correlation network analysis (WGCNA) to identify modules of genes that were co-expressed among samples in our cohort (65). We used the R package tximport (66) to import RSEM-derived gene expression counts, filtered genes for >10 counts/replicate in at least one sample, and used DESeq2 (67) to generate variance-stabilized expression counts for each gene. To detect co-expression patterns specific to deletion carriers, we used ComBat (68) within sva v.3.12 to perform batch correction with family as a covariate. We detected 35 co-expression modules in our samples using WGCNA v.1.69 (65), with the following parameters: power threshold = 8, signed hybrid network, unsigned topological overlap matrix, bi-weight mid-correlation, module size = 30-30000, and merge cut height = 0.25. Two modules showed strong sex-specific gene expression and were excluded from further analysis. The average gene expression values in each module were compared between carriers and noncarriers using two-tailed t-tests, and genes in each of these modules were tested for enrichment of Gene Ontology terms using goseq.

### Integration of gene expression and genomic variant data

We calculated the effect size of different classes of rare “second-hit” variants towards gene expression changes, stratified by sample type and family-specific patterns. We compared 25 classes of rare variants identified from WGS data (**Figure S2**; **Table S2**) towards differentially expressed genes in family trios as well as outlier expression genes in all individuals. For all comparisons, we calculated odds ratios and 95% confidence intervals for each variant class towards changes in expression using Fisher’s exact and Wald tests, respectively; uncorrected p-values and Benjamini-Hochberg corrected FDR values were reported for each comparison (**Table S5**). We note that we considered each variant class independently, so that dysregulated genes with multiple types of disrupting variants were counted within multiple variant classes. For the differential expression analysis, we first assessed variants in the 13 trios with carrier offspring for genes with differential expression (**Table S1**), and then determined the effects of variants in carrier children (n=9 trios) inherited from carrier or noncarrier parents towards expression changes shared with the same parent. For the outlier expression analysis, we first assessed variants for outlier expression genes in all individuals. We then stratified these data by sample type (carrier child, carrier parent, and noncarrier parent), and compared variants in carrier children that were inherited from their carrier or noncarrier parent. To identify synergistic effects between the 16p12.1 deletion and “second-hit” variants, we identified a subset of genes with outlier expression in deletion carriers that were also differentially expressed in the global comparison of carriers and noncarriers, and then identified those genes which also had “second-hit” variants inherited from the noncarrier parent.

### Alternative splicing analysis

To assess alternative splicing events from RNA-sequencing data, we used DESeq2 (67) to detect differential expression of isoforms. After importing isoform-level expression counts from RSEM using tximport (66), we filtered for genes with >2 counts across all samples, and performed pairwise comparisons between carrier offspring and their parents in the 13 trios listed in **Table S1**, plus the four trios with noncarrier children for comparisons. We then repeated the DESeq2 analysis for gene expression counts, and only included differentially expressed isoforms within genes that did not show overall differential expression, to specifically account for isoform changes due to alternative splicing. Similar to the family-based differential expression analysis, we assigned family-specific patterns to each alternative splicing event observed in offspring based on the pairwise comparisons to each parent. We further compared alternate isoforms identified by DESeq2 to those in GTEx LCL data (44) to identify unique isoforms in our cohort. Finally, we integrated these data with 12 classes of putative splice-site disrupting variants identified from WGS data, and calculated odds ratios as described above.

### Allele-specific expression analysis

We used the phASER v.1.1.1 (69) pipeline to identify allele-specific expression events in our cohort. We first used whatshap v.0.18 (70) to perform read-backed and pedigree-informed phasing of our WGS samples, and then merged the three replicate BAM files of aligned RNA-Seq reads for each sample together using Samtools, We then used phASER, which uses phased WGS data to infer phasing of RNA-Seq samples, to phase the RNA-Seq alignments and to count the number of reads per haplotype block. We ran phASER with the parameters --mapq 255 and - -baseq 10, and used the recommended blacklist to remove HLA genes. Finally, we quantified log-fold changes for allelic counts in each protein-coding gene with >10 read counts using phASER Gene AE, and identified ASE for genes with FDR>0.05 using binomial tests with Benjamini-Hochberg correction. For each identified ASE event, we examined the over-expressed haplotype for presence of a deleterious rare coding variant identified from WGS, which would potentially indicate pathogenic effects of the ASE event. Finally, we determined family-specific patterns of ASE genes based on the presence of ASE in parents of offspring.

### eQTL discovery and analysis

We used QTLTools v.1.2 (71) to identify eQTLs in our cohort. Because we had three replicates per participant, we first calculated the median TPM values for all transcripts in an individual. Genes were filtered for >0.1 median TPM in more than 50% of samples. Principal components for gene expression (from RNA-Seq data) and genotype (from whatshap-phased WGS data) were then computed using QTLtools. The top three genotype and the top two gene expression principal components were used as covariates for the linear model, in addition to three explicit covariates (family, sex, and carrier status). QTLtools cis-permutation tests (n=1000 replicates) were then used to discover eGenes, or genes whose expression are significantly correlated with eQTLs, and associated variants in our samples. We performed multiple testing correction with the QTLtools script runFDR_cis.R. Finally, we annotated significant eQTL variants (FDR<0.05) associated with protein-coding genes for presence in GTEx LCL data, genomic location, population frequency, and biological functions using the WGS Annovar-based pipeline (29).

### Brain-specific network analysis

We assessed the connectivity patterns of genes with “second-hit” variants and changes in expression in the context of a brain-specific interaction network. The network contains brain-specific pairwise interactions for 14,763 genes expressed in the brain, of which 11,978 (81.1%) are also expressed in the LCL samples. This network was previously built using a Bayesian classifier trained on hundreds of gene co-expression, protein-protein interaction, and regulatory-sequence datasets, in order to predict the likelihood of interactions between any two pairs of brain-expressed genes (72, 73). To create a network containing only the highest probability predicted gene interactions, we extracted all pairs of genes with weighted probabilities >2.0, representing the top ∼0.5% of pairwise interactions (217,975,718 pairs of genes). We then calculated the weighted shortest path lengths for all pairs of genes in the network, using the inverse of the probabilities as weights for each edge. Finally, we created sub-networks that contained genes with “second-hit” protein-coding variants (loss-of-function or LOF, missense, splice-site, exonic STR, or encapsulated deletion or duplication) or expression changes (differential expression, outlier expression, alternative splicing, ASE, or eQTL minor allele) for each carrier offspring from the 13 trios (**Table S1**). For each trio, we calculated the average shortest paths between all pairs of genes with expression changes and genes with “second-hit” coding variants, and then compared these distances to average shortest paths calculated from 100 permuted network replicates, where genes were randomly reassigned to different nodes in networks with otherwise identical topology. Network analysis was performed using the NetworkX package in Python (74).

### Statistical analysis

All genomic and statistical analyses were conducted using either Python v.3.7.3, with packages numpy v.1.16.2 (75), scipy v.1.1.0 (76), and pandas v.1.0.0 (77), or using R v.3.5.1 (R Foundation for Statistical Computing, Vienna, Austria). Details of all statistical tests, including summary statistics, test statistics, odds ratios, confidence intervals, p-values, and Benjamini-Hochberg corrected FDR values, are provided in **Table S5**.

## RESULTS

### The 16p12.1 deletion leads to pervasive disruption of genes involved in neurodevelopment

We performed RNA-sequencing on LCL samples from 19 deletion carriers and 13 noncarriers from five large families with multiple affected members (**Figure 1; Figure S1; Table S1**), and identified 1,569 transcripts that were differentially expressed (FDR<0.05) in deletion carriers compared with noncarriers (**Figures 2A-B**; **Table S4**). Application of additional corrections for relatedness among the samples (55) (see Methods) yielded 1,044 differentially expressed transcripts, of which 840 (80.5%) were also identified in the main analysis (**Figure S7A; Table S4**). We first confirmed that each of the seven protein-coding genes in the deletion region were downregulated in deletion carriers (**Figure 2C**). Interestingly, 13 protein-coding genes adjacent to the 16p12.1 region (between chromosomal bands 16p11.2 and 16p12.3) also showed differential gene expression in carriers, 10 of which were under-expressed in the deletion carriers. *For example*, two genes within the autism-associated 16p11.2 region, *SEZ6L2* and *DOC2A*, as well as the febrile seizure-associated gene *STX1B* (78), were downregulated in carriers of the 16p12.1 deletion. As none of the carriers harbored an atypical deletion, it is possible that these adjacent genes could be affected by disruption of regulatory elements located within the deletion region. In fact, three downregulated genes adjacent to the deletion, *DNAH3*, *OTOA*, and *NPIPB4*, exhibited chromatin interactions with enhancer elements within the deletion region, detected using published Hi-C data of LCL samples (79) (**Figure 2C**).

**Figure 1.**
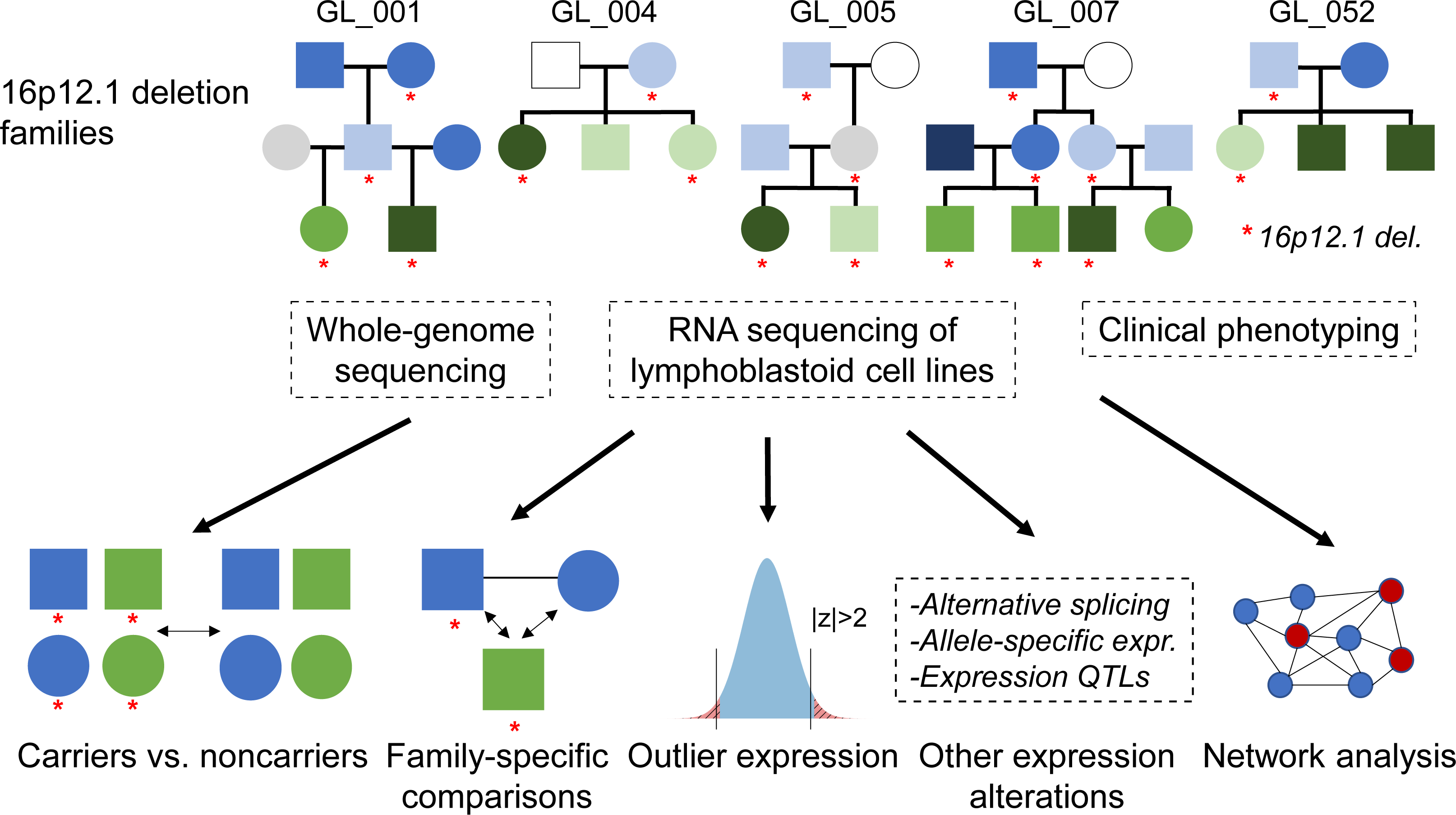
Overview of experimental design. We performed whole genome sequencing, RNA sequencing, and clinical phenotyping on five large families (32 total individuals) with the 16p12.1 deletion, indicated with red asterisks in the pedigrees. Children (green) and adults (blue) in the pedigrees are shaded by phenotypic severity score, with white indicating no clinical features, lighter shades indicating mild features (child de Vries score of 1-4; adult score of 1-2 features), medium shades indicating moderate features (child de Vries score of 5-8; adult score of 3-4 features), darker shades indicating severe features (child de Vries score of 9-13; adult score of 5-6 features), and grey indicating no phenotypic data available. Phenotypic severity scores are described in the Methods and are listed for each person in **Table S1**. We then performed multiple analyses to assess the role of the deletion and rare “second-hit” variants towards the observed transcriptomic changes and developmental phenotypes, including differential expression between carriers and noncarriers of the deletion, differential expression between parents and carrier offspring in 13 trios from the five families, outlier gene expression among all individuals, identification of additional transcriptomic alterations such as alternative splicing and allele-specific expression, and gene interaction patterns in the context of a brain-specific network.

**Figure 2.**
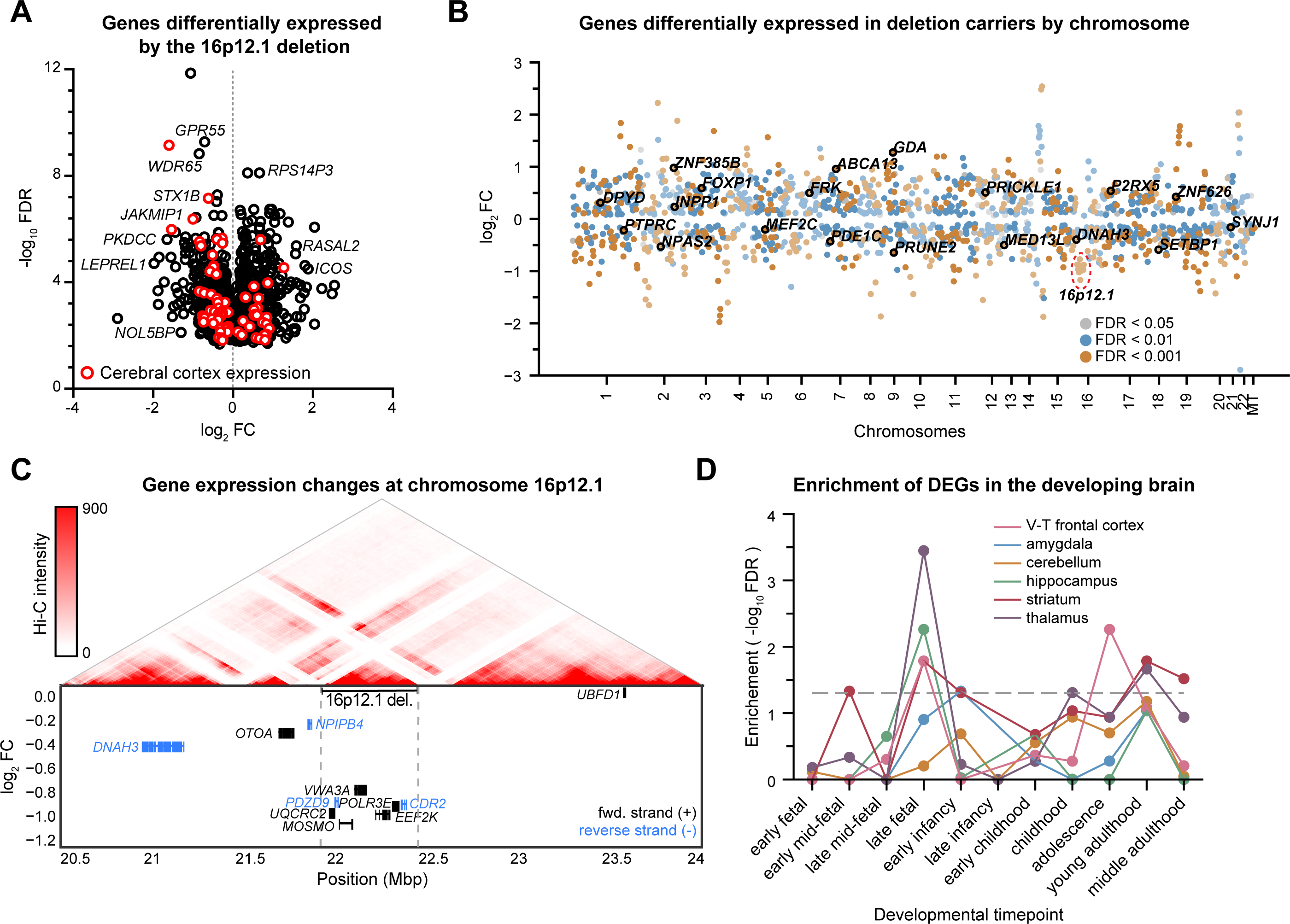
Differential expression of genes between carriers and noncarriers of the 16p12.1 deletion. **(A)** Volcano plot showing downstream (non-16p12.1 deletion) genes differentially expressed (FDR<0.05) between carriers (n=19) and noncarriers (n=13) of the deletion. Red circles indicate genes preferentially expressed in GTEx cerebral cortex tissues. **(B)** Scatter plot showing all genes differentially expressed between carriers and noncarriers of the deletion by chromosome, excluding genes on sex chromosomes. Genes are colored by FDR of differential expression. Labeled genes indicate candidate autism genes with differential expression. **(C)** Expression changes and chromatin connectivity of genes within the 16p12.1 region. The top plot shows pairwise chromatin interactions within the 3.5 Mbp 16p12.1 region, with red lines representing stronger Hi-C intensity, while the bottom plot shows log2-fold change of expression in deletion carriers of genes within and adjacent to the 16p12.1 deletion. The Hi-C data is from previously reported Hi-C experiments of LCL samples (79), and the heatmap was generated using the 3D Genome Browser (106). **(D)** Line plot shows enrichment (log10 FDR) of differentially expressed genes in deletion carriers for genes preferentially expressed in six select BrainSpan tissues across 11 developmental timepoints.

We found that differentially expressed genes in deletion carriers were enriched (FDR<0.05) for multiple biological functions, including biological adhesion and cell proliferation regulation for relatedness-corrected genes, and signal transduction and locomotion for genes without relatedness correction (**Table S4**). Additionally, we observed an enrichment (FDR=0.015) for candidate autism genes (50), including *FOXP1*, *CUL7*, *ANK3*, and *EP300*, among the differentially-expressed genes (**Figure 2B**; **Table S4**). Parametric Analysis of Gene Set Expression (PAGE) showed that genes related to neuronal and muscular growth functions were significantly upregulated in deletion carriers (FDR<0.05), while genes involved in behavioral responses and learning were downregulated (**Figure S7B; Table S6**). Weighted-gene correlation network analysis similarly identified several modules of genes with significant expression changes in deletion carriers (p<0.05, two-tailed t-test), including downregulated genes enriched for cell signaling and adhesion, and upregulated genes enriched for neurogenesis, nervous system development, and MAPK and Notch signaling (**Figure S8**; **Table S7**). Differentially expressed genes in deletion carriers were further enriched (FDR<0.05) for genes preferentially expressed in the hippocampus and basal ganglia of the adult brain (44) (**Figure S7C; Table S4**), as well as in the striatum, thalamus, and frontal cortex during late fetal and adolescent/young adulthood timepoints (61), which are critical transition periods for expression of neurodevelopmental genes (80–82) (**Figure 2D**; **Table S4**). Overall, our data suggest that the 16p12.1 deletion leads to pervasive transcriptomic changes across multiple biological and neuronal processes in the developing brain. We note that because these results are based on expression data from LCL samples, they should be followed up in neuronal models to delineate any tissue-specific differences in gene expression.

### Family-specific patterns of gene expression changes influence developmental phenotypes

We next investigated how gene expression patterns segregated within 13 complete trios with carrier offspring extracted from the five families, including carrier children compared to their parents as well as carrier parents compared with grandparents (**Table S1**). For each trio, we identified differentially expressed genes for offspring-carrier parent and offspring-noncarrier parent pairs (see Methods), and found an average of 5,323 total gene expression changes in offspring compared to their parents (**Tables S8, S9**). We then overlapped the two sets of differentially expressed genes to categorize expression changes based on their family-specific pattern (**Figure S5A**). We found no significant differences (p=0.735, two-tailed paired Mann-Whitney test) in the proportion of differentially expressed genes in offspring that were shared with either the carrier (avg. 2,223 genes/offspring) or noncarrier parent (avg. 1,908 genes/offspring; **Figure 3A**). This may suggest that “second-hit” variants from the noncarrier parent could contribute equally to gene expression changes, and therefore to disease pathogenicity, as the combined effects of the deletion and any “second-hit” variants from the carrier parent, an observation that corresponds with our recent findings of increased burden of “second-hits” transmitted to the child from noncarrier parents (16) (**Figure S5B**). However, we also note that this study may be under-powered to detect smaller differences in the proportion of gene expression changes shared between offspring and their carrier and non-carrier parents. Interestingly, we also observed an average of 1,192 genes/offspring that were differentially expressed compared with both parents (**Figure 3A**), such as *SHANK2*, *FOXP1*, and *CACNA1D*. These expression changes potentially represent effects of *de novo* variants or combinatorial effects of variants inherited from both parents, which could explain the increased phenotypic severity observed in the carrier children. However, the trends in expression patterns widely varied across families, which in some cases could be explained by family history of neuropsychiatric disease (**Figure S5C**). For example, we found that children within family GL_004, whose parents were unaffected or presented with mild depression, had the lowest number of gene expression changes among any carrier children in the cohort. Meanwhile, children in families GL_001 and GL_052, whose carrier parents manifested multiple overt cognitive and neuropsychiatric features, had higher proportions of expression changes shared with their carrier parents compared to their noncarrier parents (**Figure S5C**).

**Figure 3.**
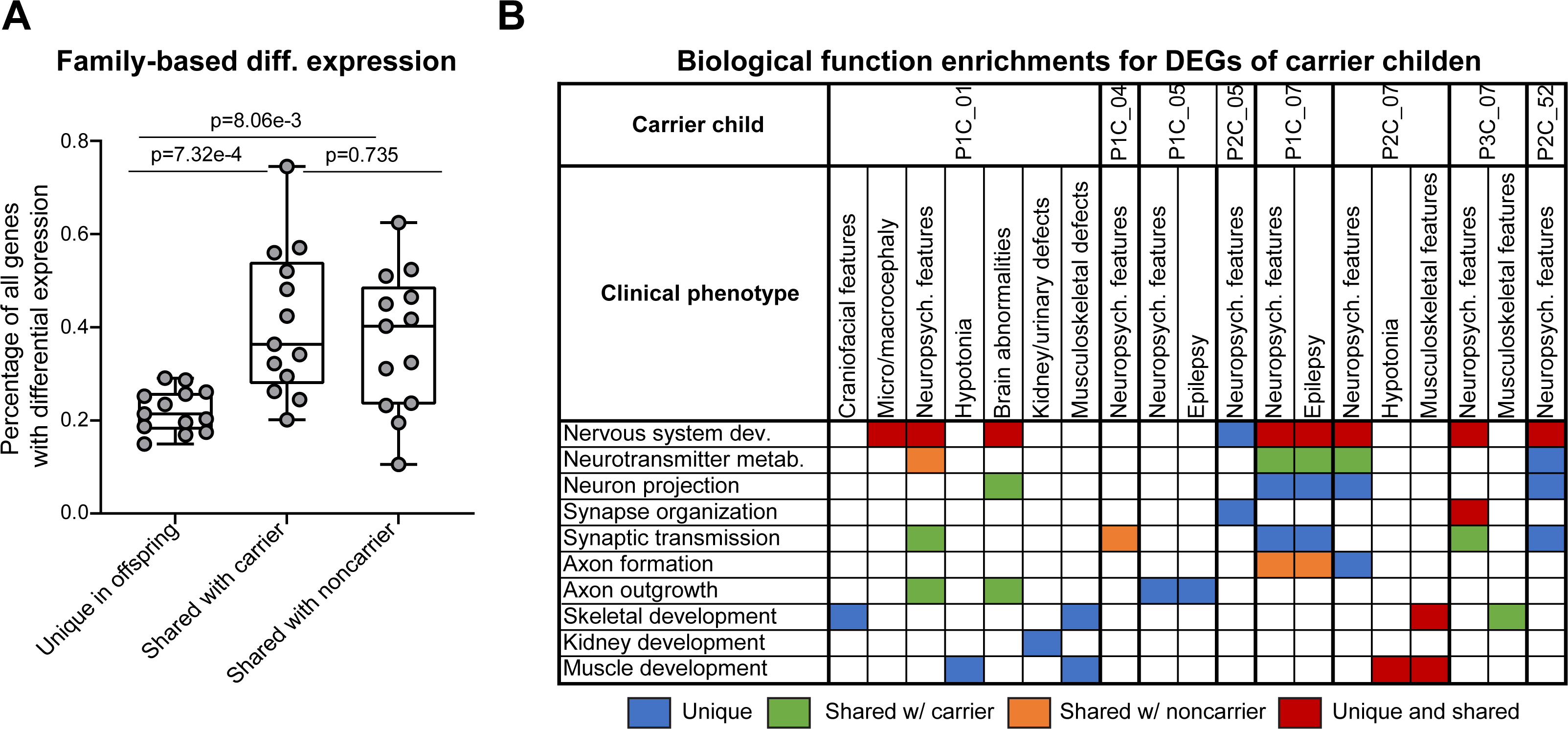
Differential expression of genes between offspring and carrier and noncarrier parents. **(A)** Boxplot shows the proportion of differentially expressed genes in carrier offspring of 13 trios (**Table S1**) that were either unique to the offspring or shared with their carrier or noncarrier parents (*p<0.05, two-tailed paired Mann-Whitney test). Boxplot indicates median (center line), 25th and 75th percentiles (bounds of box), and minimum and maximum (whiskers). **(B)** Table shows observed clinical features in eight carrier children with overt developmental phenotypes, as well as enrichments (FDR<0.05) of differentially expressed genes in each carrier child for biological functions related to each clinical feature. Cells are colored according to the family-specific patterns (uniquely observed or shared with a parent) of differentially expressed genes for each enriched biological process.

We next assessed the dysregulated biological functions in each trio (**Figure S9**; **Table S9**), and found that unique or shared differentially expressed genes in carrier children were enriched for biological processes (FDR<0.05) that could be related to 33 out of 41 (80.5%) developmental phenotypes observed in the affected children (**Figure 3B**). For example, shared gene expression changes for carrier child P1C_01 in family GL_001 were enriched for nervous system development, neurotransmitter metabolism, neuron projection, and synaptic transmission functions, while their unique expression changes were enriched for genes involved in skeletal and muscular development. The shared changes in neuronal genes could contribute to the ID/DD and speech delay phenotypes observed in the child, as both parents also had several psychiatric features, while the unique changes in developmental genes could be related to hypotonia, growth delay, and craniofacial features uniquely observed in the child. Overall, these results suggest that expression changes of neurodevelopmental-related genes could account for phenotypic differences among carriers of the 16p12.1 deletion.

### “Second-hit” variants and the 16p12.1 deletion show synergistic effects towards gene expression

We next investigated whether changes in gene expression could be attributed to “second-hits”, or rare genetic modifiers elsewhere in the genome. Rare variants disrupting protein-coding regions and nearby regulatory elements have been previously linked to gene expression changes in both control populations (23,83–85) and disease cohorts, where causal genes may be missed by DNA sequencing methods (18,20–22). We hypothesized that “second-hits” by themselves or in combination with the deletion could contribute to the observed gene expression changes in affected children. We therefore identified 25 classes of rare gene-disruptive “second-hit” variants from WGS data for each individual (**Figure S2**; **Tables S9, S10**), including SNVs and indels in coding and non-coding regulatory regions (UTRs, introns, and putative promoter, enhancer, and silencer elements within 50 kbp of a gene) with Phred-like CADD scores>10 (34), and CNVs and short tandem repeats (STRs) that spanned gene-coding regions. We then calculated the likelihood that these “second-hit” variants are associated with changes in expression of a proximal gene, as determined by either differential expression analysis between carrier offspring and their parents in the 13 trios, or outlier expression analysis among all individuals in the cohort (18, 83) (see Methods). While family-based differential expression analysis detects all expression changes between affected children and their parents (86), including those due to the downstream effects of the deletion, outlier analysis more robustly identifies specific effects of “second-hits” towards larger changes in expression, including synergistic effects in combination with the deletion. Overall, we observed an average of 285 outlier genes (|z-score| >2) per individual, including candidate neurodevelopmental genes (49) such as *CTNNB1*, *FOXG1*, *DISC1*, and *ZNF804A* (**Figure S6**; **Tables S8, S11**). We found that 10.8% of outlier genes (avg. 31/286 per person) and 11.2% of differentially expressed genes (avg. 310/2,774 per carrier offspring) were potentially disrupted by a rare coding or non-coding variant (**Table S10**). Altered expression of genes without such variants could be due to several factors, such as common variants, DNA methylation events, downstream effects of other dysregulated genes, or environmental factors.

In agreement with previous studies (23, 83), we found that genes with outlier expression were significantly enriched after Benjamini-Hochberg correction (FDR<0.05, Fisher’s exact test) for 5/25 classes of rare variants that directly affected gene-coding regions, including loss-of-function (LOF), missense, and splice-site SNVs, and 5’ UTR overhanging and gene-encapsulating duplications (**Figure 4A**). Similarly, we found that 10/25 variant classes were significantly associated with differentially expressed genes in carrier offspring for the 13 trios (FDR<0.05, Fisher’s exact test), including coding missense SNVs, duplications overhanging the 5’ UTR, and encapsulated deletions (**Figure S10A**). We further found that outlier genes had higher burdens of rare variants in aggregate (p=1.01×10^-3^, one-tailed t-test) and for 7/25 individual classes compared with non-outlier genes (p<0.05, one-tailed t-test), in particular loss-of-function variants (FDR=1.73×10^-3^) and encapsulated duplications (FDR=5.43×10^-3^), which passed multiple-testing correction (**Figures S11A-B**). Interestingly, we also found that outlier genes that were intolerant to variation (pLI score >0.9 or RVIS percentile <20; p<3.26×10^-4^, two-tailed t-test) or preferentially expressed in the brain (p=0.011) had a higher burden of nearby rare variants than other outlier genes (**Figure S11C**). We next assessed the effect size of “second-hits” towards outlier gene expression among carriers and noncarriers of the deletion, and found enrichments of LOF variants (p=4.96×10^-6^; FDR=6.20×10^-5^), 5’ UTR overhanging duplications (p=0.017; FDR=0.108), and 5’ UTR-disrupting SNVs (p=7.37×10^-3^; FDR=0.058) towards outlier expression in carrier children but not in carrier parents (**Figure 4B**). We observed similar findings among differentially expressed genes, where missense SNVs (FDR= 4.23×10^-5^), upstream SNVs (FDR=7.00×10^-4^), encapsulated (FDR=0.039) and interstitial (FDR=0.031) deletions, and 5’ UTR overhanging duplications (FDR=0.045) were only likely to alter gene expression in carrier children (**Figure S10B**). Notably, LOF variants (p=1.67×10^-5^, FDR=2.09×10^-4^) and 5’ UTR overhanging duplications (p=0.018; FDR=0.090) were also enriched for outlier expression in noncarrier parents, suggesting that these classes of “second-hit” variants were more likely to be deleterious in carrier children when inherited from noncarrier parents (**Figure 4B**). In fact, we observed several classes of “second-hit” variants in carrier children, including 5’ UTR overhanging duplications (p=1.54×10^-3^, FDR=0.025), gene-encapsulating (p=0.025; FDR=0.125) and 3’ UTR overhanging deletions (p=0.025; FDR=0.125), and missense (p=0.023; FDR=0.125) and upstream SNVs (p=0.032; FDR=0.133), that were enriched for outlier expression when inherited from the noncarrier parent but not from the carrier parent (**Figure 4C**). Similarly, we found that LOF variants (FDR=5.05×10^-5^) and SNVs in upstream (FDR=6.25×10^-3^) and silencer regions (FDR=7.70×10^-4^) correlated with differential gene expression in carrier children only when inherited from the noncarrier parent (**Figure S10C**). For example, a carrier child in family GL_007, who exhibited hypotonia and muscle weakness, inherited a deleterious variant from their noncarrier parent in the 5’ UTR of *EIF2AK1*, associated with motor dysfunction (87), that potentially led to downregulation of that gene (**Figure 4D**). Overall, our findings showed that distinct classes of “second-hit” variants differentially contribute towards changes in gene expression when inherited in a complex manner from either the carrier or noncarrier parent.

**Figure 4.**
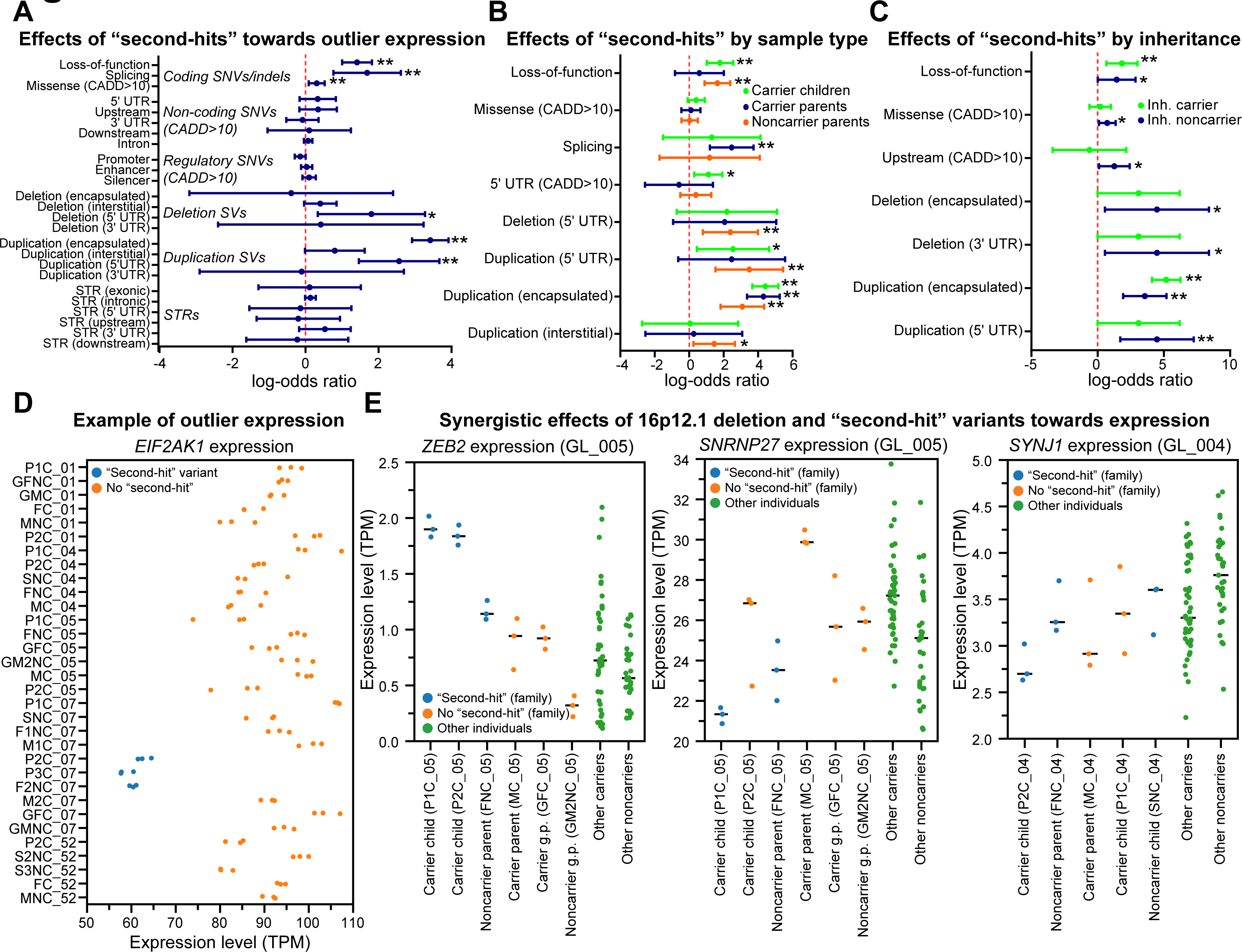
Enrichment of “second-hit” variants near genes with outlier expression. **(A)** Forest plot shows enrichment (Fisher’s exact test, **=FDR<0.05, *=uncorrected p<0.05) of genes with outlier expression in all individuals in the cohort (n=32) for rare proximal coding and non-coding variants, including single-nucleotide variants (SNVs) and insertions/deletions (indels) with CADD scores >10 (34), structural variants (SVs), and short tandem repeats (STRs). **(B)** Forest plot shows classes of “second-hit” variants with significant enrichment (Fisher’s exact test, **=FDR<0.05, *=uncorrected p<0.05) towards genes with outlier expression in carrier children (n=10), carrier parents (n=6), or noncarrier parents (n=6). **(C)** Forest plot shows classes of “second-hit” variants with significant enrichment (Fisher’s exact test, **=FDR<0.05, *=uncorrected p<0.05) towards genes with outlier expression in carrier children (n=9) that are shared with either carrier or noncarrier parents. All forest plots show log-odds ratios (dots) and 95% confidence intervals (whiskers). Odds ratios, confidence intervals, p-values, and Benjamini-Hochberg corrected FDR values for comparisons with all classes of “second-hit” variants are listed in **Table S5**. **(D)** Scatter plot shows expression values (transcripts per million, or TPM) for *EIF2AK1* in LCL replicates for all individuals (n=32). Samples in blue have outlier expression of *EIF2AK1* (z-score <-2) and carry a deleterious “second-hit” variant in the 5’ UTR of the gene. **(E)** Scatter plots show expression values (TPM) of genes with synergistic effects due to the 16p12.1 deletion and inherited “second-hit” variants. Blue circles indicate expression values for samples from carrier children and family members with rare “second-hit” variants, orange circles indicate expression values for samples from family members without the “second-hit” variant, and green circles indicate expression values of samples from other deletion carriers and noncarriers in the cohort. Black lines denote median gene expression for LCL replicates of each individual used to identify genes with outlier expression in individual deletion carriers.

We next investigated whether “second-hit” variants showed synergistic effects towards expression changes in genes also dysregulated by the 16p12.1 deletion. We found 11 instances of genes, such as *ANK3, DOCK10,* and *SLC26A1*, that were differentially expressed in all deletion carriers, showed outlier expression in an individual deletion carrier, and had a nearby variant (two coding and nine non-coding) inherited from the noncarrier parent (**Figure 4E**; **Figure S12**; **Table S12**). For example, two carrier children in family GL_005 inherited an intronic variant within *ZEB2* from their noncarrier parent, whose altered dosage is associated with Mowat-Wilson syndrome (88, 89). While both the 16p12.1 deletion and the “second-hit” variant individually corresponded with increased *ZEB2* expression, the presence of both variants in the carrier children resulted in even stronger overexpression of the gene compared to those with either individual variant (**Figure 4E**). Overexpression of *ZEB2* could contribute to the Mowat-Wilson like features observed in the carrier child P1C_05, including ID/DD, seizures, hypotonia, and digestive abnormalities. Similarly, a carrier child in GL_005 inherited a rare variant in a promoter region upstream of the mRNA splicing-associated (90) gene *SNRNP27* from their noncarrier parent. *SNRNP27* is over-expressed in deletion carriers but under-expressed in both the carrier child and the noncarrier parent, highlighting a case where a “second-hit” variant reverses an expression change caused by the deletion (**Figure 4E**). Furthermore, a carrier child in GL_004 shared an intronic variant with two noncarrier relatives in the gene *SYNJ1*, which is associated with synaptic transmission (91) and is under-expressed in carriers of the deletion. While other individuals with the same variant had normal *SYNJ1* expression, the carrier child exhibited under-expression of the gene compared to both carriers and noncarriers of the deletion, suggesting that the variant may alter *SYNJ1* expression only in the presence of the deletion (**Figure 4E**). While it is possible that other variants elsewhere in the genome could also influence expression levels of these genes, these examples highlight putative synergistic effects between the 16p12.1 deletion and “second-hit” variants towards gene expression, where the “second-hit” variants may amplify or reduce the effects of the CNV.

### A broad range of transcriptomic alterations contribute to phenotypic variability of the 16p12.1 deletion

To identify a complete spectrum of gene expression alterations in each individual, we next evaluated alternative gene splicing, allele-specific expression (ASE), and expression quantitative trait loci (eQTL) among individuals in our cohort. We first identified an average of 3,267 alternative isoforms present in carrier offspring of the 13 trios compared to their parents (**Tables S8, S13**), including for several neurodevelopmental-associated genes (49) such as *KANSL1*, *SHANK2*, and *SYNGAP1*. After categorizing splicing events by family-specific patterns, we found no differences between splicing events in offspring shared with carrier (average=1,307) or noncarrier parents (average=1,392; p=0.635, two-tailed paired Mann-Whitney test), with fewer unique changes in the offspring (average=568; p=2.44×10^-4^; **Figure S13A**). We next found enrichments for alternative splicing in genes disrupted by “second-hit” splice-site (p=7.47×10^-4^, Fisher’s exact test; FDR=2.99×10^-3^), intronic (p=9.91×10^-9^; FDR=5.95×10^-8^), or missense SNVs (p=0.012; FDR=0.036), interstitial (p=0.043; FDR=0.086) and 3’ UTR overhanging deletions (p=0.024; FDR=0.058), and intronic STRs (p=5.75×10^-10^; FDR=6.90×10^-9^) (**Figure S13B; Table S10**). We also found that intronic SNVs were more likely to disrupt splicing in carrier children if they were inherited from the noncarrier parent (p=0.034, FDR=0.204) than the carrier parent (**Figure 5A**), while intronic SNVs (FDR=6.36×10^-7^) and interstitial deletions (FDR=0.018) were more likely to lead to alternative splicing when present in carrier children than in carrier parents (**Figure S13C**). These results suggest potential correlations between classes of inherited rare variants and alternative splicing events, although changes in isoform expression can only be confidently attributed to rare variants at or near the splice-site. For example, a deleterious splice-site variant in the transcriptional activator *TADA2A* led to an alternate isoform (*TADA2A*-003) in multiple family members of GL_007 that was not observed in GTEx LCL data (**Figure 5B**). *TADA2A* is a candidate gene within the schizophrenia-associated 17q12 deletion (92), and five out of six family members with the splicing variant have schizophrenia-like clinical features (i.e. hallucinations or delusions), including four deletion carriers and one noncarrier child.

**Figure 5.**
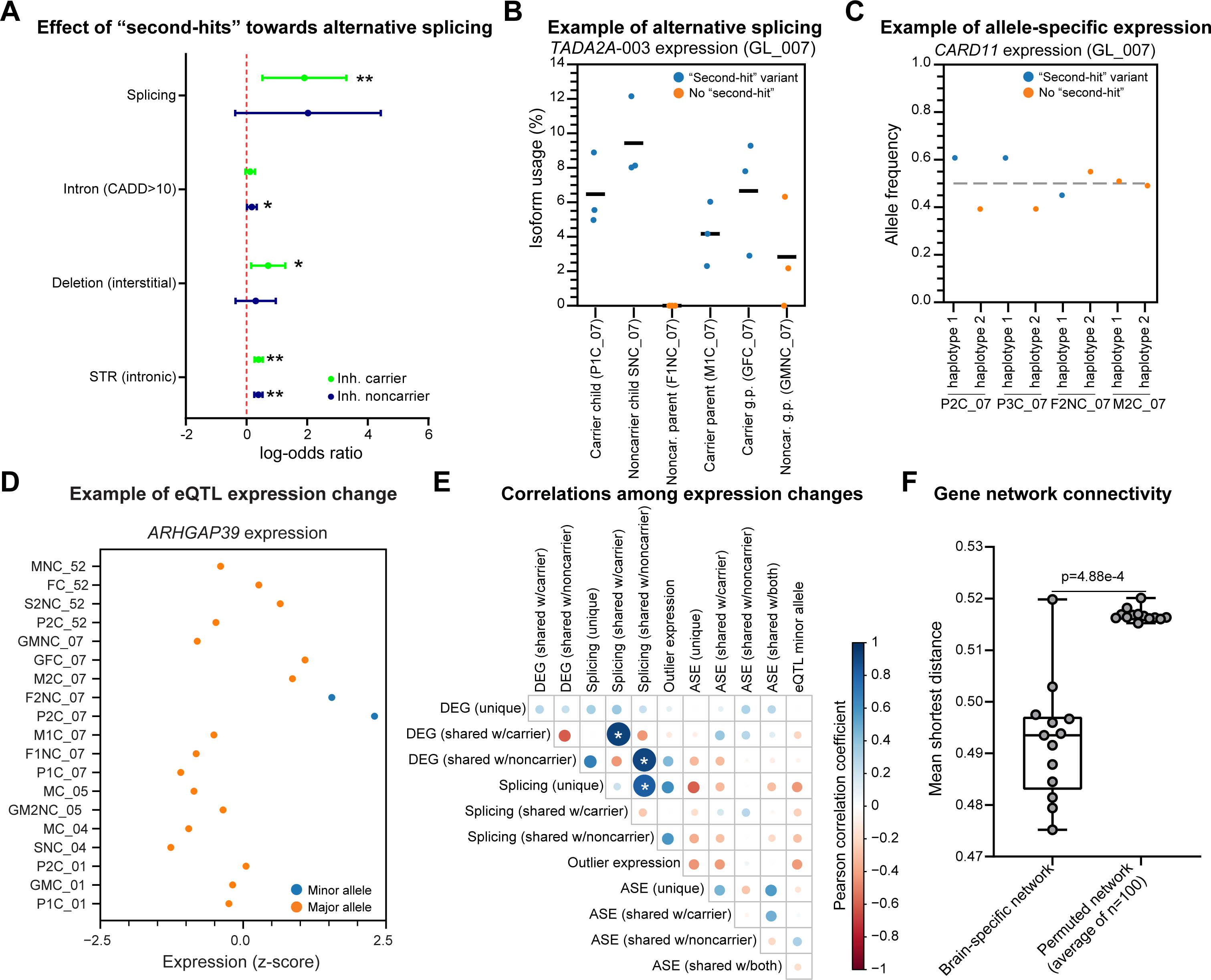
Alternative splicing, allele specific expression, eQTL, and network analysis. (A) Forest plot shows classes of rare variants with significant enrichment (Fisher’s exact test, **=FDR<0.05, *=uncorrected p<0.05) towards genes with alternative splicing in carrier children (n=9) that are shared with either carrier or noncarrier parents. Forest plot shows log-odds ratios (dots) and 95% confidence intervals (whiskers). Odds ratios, confidence intervals, p-values, and Benjamini-Hochberg corrected FDR values for comparisons with all classes of “second-hit” variants are listed in **Table S5**. (**B)** Scatter plot shows isoform usage percentage for *TADA2A-* 003 in replicates for individuals in family GL_007. Samples in blue carry a “second-hit” splice-site variant in *TADA2A* and exhibit a higher frequency of the alternative isoform. **(C)** Scatter plot shows allele frequencies for the autism-associated gene *CARD11* in carrier child P2C_07, noncarrier parent F2NC_07, and carrier parent M2C_07 in GL_007. Blue circles indicate allele frequency for haplotypes carrying a “second-hit” coding variant disrupting *CARD11*. **(D)** Scatter plot shows z-scores for expression values of *ARHGAP39* for all individuals with available genotypes for the gene. Individuals who carry the minor allele for the *ARHGAP39* eQTL (blue dots) have higher expression of the gene than the rest of the cohort (orange dots). **(E)** Plot shows correlations among the numbers of gene expression alterations in carrier offspring for the 13 trios assessed in our study. Colors and sizes of the circles are proportional to the correlation coefficients between gene expression changes, where blue indicates a positive correlation and red indicates a negative correlation. Asterisks denote significant correlations (FDR<0.05, Pearson correlation with Benjamini-Hochberg correction). **(F)** Boxplot shows the average shortest distances for carrier offspring (n=13) between pairs of genes with “second-hit” coding variants and genes with identified expression changes in a brain-specific network. Genes with expression changes were more strongly connected to genes with “second-hit” variants in the brain-specific network than the average distances for genes within 100 permuted brain-specific networks per sample (p=4.88×10^-4^, two-tailed paired Mann-Whitney test). Boxplot indicates median (center line), 25th and 75th percentiles (bounds of box), and minimum and maximum (whiskers).

Next, we identified an average of 285 genes with ASE per individual in our cohort (**Tables S8, S14**), including for the neurodevelopmental-associated genes (49) *DNMT3A*, *NSUN2*, and *HDAC8*. ASE events in the 13 trios were more likely to uniquely occur in the offspring than be shared with a parent (p=2.44×10^-4^, two-tailed paired Mann-Whitney test), in contrast to differential expression and alternative splicing events (**Figure S14A**). Genes with ASE have previously been shown to have a higher burden of nearby rare deleterious variants (83), and the pathogenicity of a gene with ASE increases with the presence of a deleterious variant on the overexpressed allele (85). In our cohort, we found five ASE events in carrier children that led to overexpression of a deleterious “second-hit” coding variant (**Figure S14B**). For example, two carrier children with autism in family GL_007 showed overexpression of a deleterious “second-hit” missense variant in the candidate autism gene (93) *CARD11,* which was inherited from their noncarrier parent (**Figure 5C**).

We further performed eQTL discovery analysis to identify variants statistically correlated with expression changes in our cohort, agnostic to variant pathogenicity or population frequency. We identified 21 eQTLs which modulated the expression of 23 eGenes, or genes whose expression is significantly correlated with an eQTL (**Figure S15A; Table S15**). Interestingly, 19/21 identified eQTLs were not present in GTEx LCL data, representing unique loci in our cohort. Carrier children showed a trend (p=0.107, two-tailed Mann-Whitney test) towards carrying a higher number of minor eQTLs alleles (average=4.3/person) than their carrier parents (average=3.2/person) (**Figure S15B; Table S8**). Furthermore, several eGenes had biological functions related to neuronal processes (94–96) (**Table S15**), including *SERPINF1, BEGAIN,* and *ARFGEF2*. For example, we identified a relatively rare eQTL (allele frequency = 0.015) for overexpression of *ARHGAP39*, a key regulator of neurogenesis and dendrite morphology associated with learning and memory (97) (**Figure 5D**). The eQTL minor allele, located in a transcription factor binding cluster, was only observed in a carrier child and their noncarrier parent within GL_007, who both presented with neuropsychiatric phenotypes.

To assess the joint contributions of each type of expression change among the individuals in our cohort, we assessed correlations between the numbers of gene expression changes assessed in our study by family-specific pattern (**Figure 5E**). We observed three significant positive correlations (FDR<0.05, Pearson correlation) between pairs of gene expression changes in each person, which often shared the same family-specific patterns. Specifically, the number of genes with differential expression strongly correlated with the number of genes with alternative splicing when shared with either the carrier parent (r=0.93, FDR=2.91×10^-4^) or non-carrier parent (r=0.91, FDR=4.52×10^-4^), while unique splicing events in the offspring correlated with splicing events shared with the non-carrier parent (r=0.83, FDR=0.011). Together, the correlations between transcriptomic alterations suggest that different types of gene expression changes could co-occur in parents and offspring, potentially due to the same inherited “second-hit” variants disrupting expression of similar genes and biological pathways, as is observed for signals in genome-wide association studies (98).

### Genes with “second-hit” variants and expression changes show strong connectivity in a brain-specific network

Finally, to determine whether associations between transcriptomic changes and “second-hit” variants in LCL samples were also relevant in the nervous system, we assessed connectivity patterns of genes with “second-hit” variants and altered gene expression using a brain-specific gene interaction network (72, 73). We generated individual networks for carrier offspring in the 13 trios, and calculated shortest distances between genes with protein-coding “second-hit” variants and genes with LCL-derived expression changes in each offspring (see Methods). We found that the average shortest distances between genes with “second-hits” and expression changes were significantly smaller in 6/13 offspring than those derived from permuted networks (FDR<0.05, one-tailed z-test with n=100 permutations). In fact, networks for offspring in aggregate had significantly smaller shortest distances (p=4.88×10^-4^, two-tailed paired Mann-Whitney test) than the shortest distances from the sets of permuted networks, where genes were randomly reassigned to different nodes in the network (**Figure 5F**). These data indicate that “second-hit” variants closely interact with genes with expression changes detected from LCL samples in a brain-specific context, suggesting a potential mechanism for how gene expression changes that underlie developmental phenotypes can be influenced by “second-hit” variants in the genome. However, these findings should be confirmed using expression data from patient-derived neuronal models of the 16p12.1 deletion, as expression changes in LCL samples may not be conserved in the nervous system.

## DISCUSSION

We previously described a two-hit model for the 16p12.1 deletion, where the presence of both the deletion and “second-hit” variants determine the phenotypic trajectory of affected children (11, 16). Here, we propose a potential mechanism for how the deletion and “second-hits” jointly interact to alter clinical phenotypes by way of the transcriptome. We found that the 16p12.1 deletion itself disrupts the expression of genes across the genome through direct effects, such as chromatin interactions, and through indirect effects, such as downstream genetic interactions (**Figure 6**). For example, chromatin interactions were observed between regions within the 16p12.1 deletion and flanking genes such as *STX1B* and *DNAH3*, and 1,493 genes outside of chromosome 16 were also dysregulated in deletion carriers. The identification of flanking genes downregulated by the deletion is in line with similar findings for the 16p11.2, 1q21.1, and 22q11.2 deletion disorders (19, 99). Each of these CNVs exhibited altered gene expression in adjacent regions that is putatively mediated by chromatin interactions, highlighting the importance of considering the three-dimensional structure of the genome to elucidate CNV pathogenicity. Similarly, we found that “second-hits” disrupt gene expression through both direct and indirect mechanisms. Genes with nearby “second-hit” variants were more likely to exhibit outlier expression and alternative splicing, and genes with “second-hits” were more closely connected to genes with expression changes in a brain-specific network than random sets of genes in permuted networks. In fact, we observed 11 examples of combined effects of the deletion and “second-hit” variants towards expression in our cohort, including the candidate neurodevelopmental genes *SYNJ1* and *ZEB2*. These synergistic effects towards gene expression are similar to those previously observed for eQTLs (100) and HLA alleles (101), except that these effects are potentially due to the combined effects of rare deleterious variants. We note that only a subset (∼11%) of genes with altered expression in our cohort harbored deleterious “second-hit” variants that could affect expression. It is likely that the downstream effects of both the deletion and “second-hit” variants could be responsible for a larger proportion of gene expression changes, along with other common variants and environmental factors. Thus, our results suggest that the 16p12.1 deletion and the “second-hit” variants interact with each other in a complex manner to mold the shape of the transcriptome, resulting in strong dysregulation of developmental genes and contributing to neuropsychiatric features in 16p12.1 deletion carriers. Results from our study align with previous studies, which found that rare variants of different classes have varying effect sizes towards gene expression (23, 83). Our study extends this paradigm by identifying classes of rare “second-hit” variants whose contributions to gene expression changes differ by inheritance pattern. We found that high-effect variants, such as whole-gene duplications, cause expression changes regardless of parent-of-origin, and splice-site variants lead to changes in isoform expression independently of inheritance. In contrast, lower-effect variants, including missense, silencer, and upstream SNVs, were more strongly associated with gene expression changes when inherited from the noncarrier parent than the carrier parent. These findings indicate that noncarrier parents are more likely to pass gene expression-altering “second-hit” variants down to their carrier children, potentially accounting for more severe phenotypic manifestations in children with the deletion compared with their carrier parents (**Figure 6**). One potential explanation for why carrier children receive a higher number of deleterious variants from their non-carrier parent is assortative mating among their parents, as 8/8 carrier parents and 7/9 non-carrier parents in our cohort manifested at least mild neuropsychiatric features. Assortative mating has been extensively observed among individuals with neurodevelopmental or psychiatric disorders (102, 103), in particular autism (104), suggesting its relevance towards phenotypic variability among deletion carriers on our cohort. Future family-based transcriptome studies with larger sample sizes may be able to pinpoint specific rare variants within dysregulated genes that are associated with distinct phenotypes in the carrier children.

**Figure 6.**
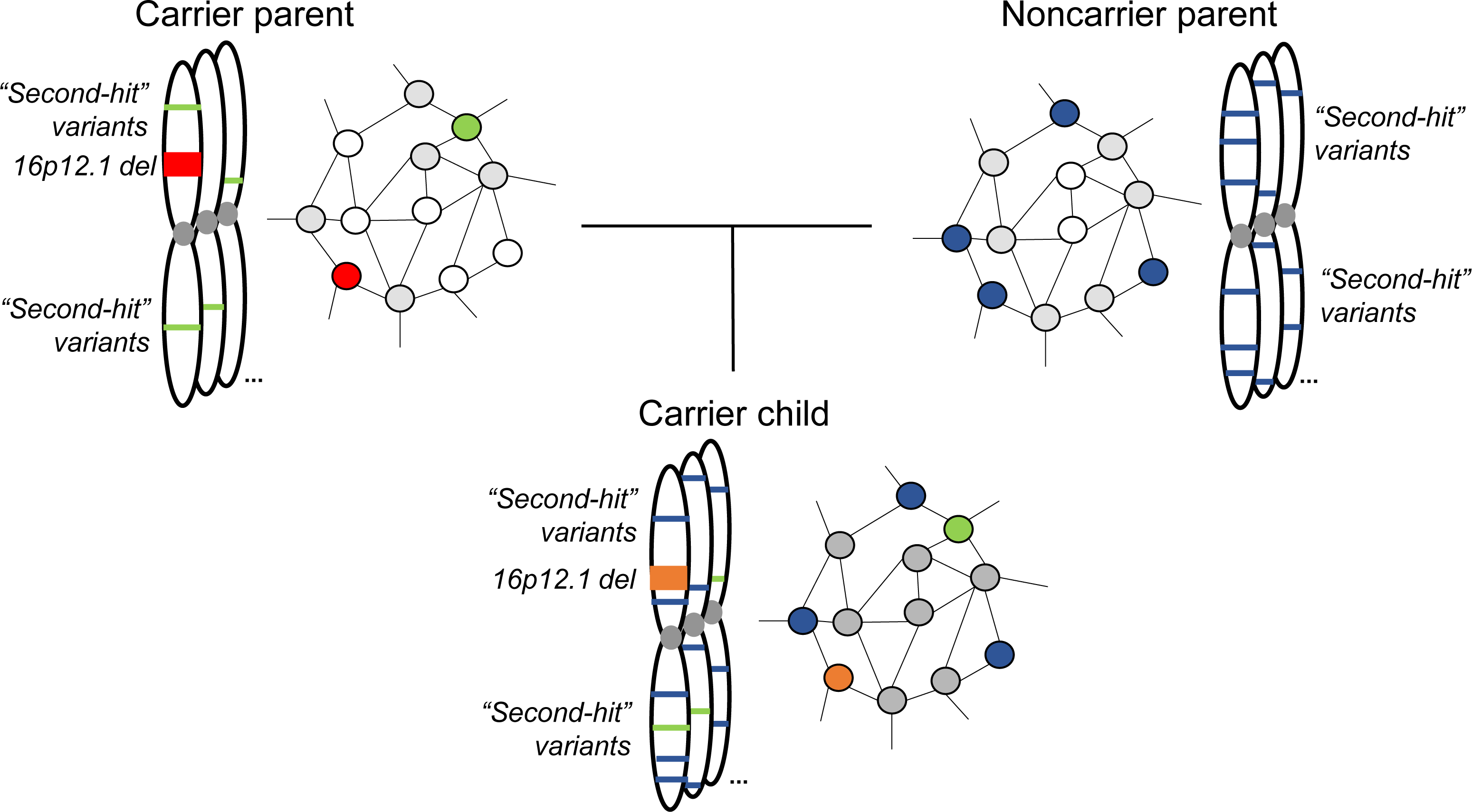
Genetic and transcriptomic mechanisms for phenotypic variability in 16p12.1 deletion families. Affected children inherit the 16p12.1 deletion (red) and a smaller number of rare “second-hit” variants (green) from a carrier parent, and a larger number of “second-hit” variants from the noncarrier parent (blue). Altered expression of genes due to these “second-hit” variants affects nearby downstream connected genes in an interaction network (grey), causing additional transcriptomic perturbation. Because of this, carrier children have numerous gene expression changes compared with their carrier parents, including genes showing synergistic effects of the deletion and “second-hit” variants (orange), potentially accounting for more severe developmental phenotypes observed in the children.

We also identified putative biological and developmental pathways disrupted by both the deletion and “second-hit” variants. For example, we found that genes differentially expressed by the deletion were preferentially expressed in multiple brain tissues during development, and were enriched for core signaling and developmental pathways. In fact, knockdown of individual homologs of 16p12.1 genes in *Drosophila melanogaster* models showed neuronal phenotypes and transcriptome disruptions, suggesting that the individual effects of multiple genes in the deletion sensitize the genome for neuropsychiatric phenotypes (105). Interestingly, we found several examples of biological functions and mechanisms that were simultaneously dysregulated by both the deletion and “second-hit” variants in the carrier children. For example, most carrier children shared differential expression in genes enriched for nervous system development, cell adhesion, signaling, and locomotion with both their carrier and noncarrier parents. These results provide insights into how the deletion and “second-hit” variants synergistically dysregulate genes and pathways related to development, ultimately contributing towards a wide range of developmental phenotypes observed in children with the deletion.

Some limitations can be noted in the context of our study. *First*, we investigated gene expression changes within patient-derived LCL samples, which have reduced relevance for brain expression. However, over 80% of genes expressed in GTEx brain samples, as well as over 70% of neurodevelopmental disease genes, were expressed in our LCL samples (**Figures S4A-B**). Nevertheless, repeating the study in tissues that are implicated in neurodevelopmental disorders, potentially using patient-derived reprogrammed neuronal progenitor cells, would verify the associations between variants, expression changes, biological functions, and clinical features. *Second*, we have a relatively small cohort of 32 individuals within five families. It would be useful to determine whether the identified associations are strengthened in a larger cohort, especially those that did not pass multiple-testing corrections. Phenotypically more diverse cohorts would also allow for performing additional correlations between gene expression changes and specific clinical features, such as whether more outlier genes are present among families with stronger histories of neuropsychiatric disease.

## CONCLUSIONS

Overall, our work supports a model for complex disorders, where combinatorial effects of multiple variants with different effect sizes affect expression of genes in developmental pathways, which further influence the expressivity of clinical features. These results exemplify that family-based transcriptome studies, similar to family-based genome studies, can help explain changes in phenotypes from parents to children and between siblings, especially in complex disorders with a high degree of intra- and inter-familial variability.

## LIST OF ABBREVIATIONS

ASE: allele-specific expression
CNV: copy-number variant
ID/DD: intellectual disability/developmental delay
LCL: lymphoblastoid cell line
LOF: loss-of-function
PAGE: parametric analysis of gene set enrichment
RIN: RNA integrity number
SNV: single-nucleotide variant
STR: short tandem repeat
TPM: transcripts per million
TSS: transcription start site
UTR: untranslated region
WGCNA: weighted gene correlation network analysis
WGS: whole genome sequencing

## DECLARATIONS

### Ethics approval and consent to participate

Carrier children and their family members provided informed consent according to a protocol reviewed and approved by the Pennsylvania State University Institutional Review Board (IRB #STUDY00000278).

### Consent for publication

Not applicable.

### Availability of data and materials

Patient-derived LCL samples generated in this study are available at the NIGMS Human Genetic Cell Repository at the Coriell Institute (https://www.coriell.org/1/NIGMS). Accession numbers for LCL samples are provided in **Table S3**. Whole genome sequencing, SNP microarray, and RNA-sequencing data generated in this study are available at NCBI dbGaP study accession phs002450 (http://www.ncbi.nlm.nih.gov/projects/gap/cgi-bin/study.cgi?study_id=phs002450.v1.p1) and BioProject accession number PRJNA734670. All other data generated or analyzed during this study are included in this article and its supplementary information files. All code generated for this project, including pipelines for running bioinformatic software and custom analysis scripts, are available at https://github.com/girirajanlab/16p12_RNAseq_project and https://github.com/girirajanlab/16p12_WGS_project.

### Authors’ contributions

MJ, AT, and SG conceived the project. MJ and AT performed all analyses in the manuscript, generated the plots and images, and wrote and revised the manuscript. LP recruited the families, obtained and assessed clinical phenotypes, and isolated DNA and RNA for sequencing. CS assisted with collection of patient clinical information and bioinformatic pipelines to identify WGS variants. MD assisted with extraction of RNA from LCL samples. EH assisted with collection of patient clinical information. AK provided the brain-specific network and assisted with the network and WGCNA analyses. SG supervised the research, recruited the families, assisted with collection of patient clinical information, and wrote and revised the manuscript. All authors read and approved the final draft of the manuscript.

## Acknowledgements

We are grateful to all members of the five families who participated in this study. We also thank Sherryann Wert, Melissa Berkowitz, and Tara Schmidlen (Coriell Institute) for their assistance in the generation of stable LCL lines; Craig Praul and Yuka Imamura (Penn State Genomics Core Facility) for assistance with designing the RNA-sequencing strategy; Casey Brown (UPenn), Istvan Albert (Penn State), and Qunhua Li (Penn State) for assistance with statistical analysis of transcriptome data; and Jesse Gillis (CSHL), Dajiang Liu (Penn State), and members of the Girirajan lab for useful discussions and critical reading of the manuscript. We appreciate access to data from the Genotype-Tissue Expression (GTEx) Project, which was supported by the Common Fund of the Office of the Director of the National Institutes of Health, and by NCI, NHGRI, NHLBI, NIDA, NIMH, and NINDS. The data used for the analyses described in this manuscript were obtained from the GTEx Portal in January 2020.

## ADDITIONAL FILES

**Additional file 1:** Fifteen supporting Figures S1-S15. A figure caption for each is given within the file (Format: PDF).

**Additional files 2-16:** Individual files for supporting Tables S1-S15 (Format: Excel). Table captions are as follows:

**Table S1.** This file lists 32 individuals in the five 16p12.1 deletion families by family relationship, sex, deletion carrier status, and observed developmental or neuropsychiatric phenotypes, including modified de Vries scores for children and adult phenotypic severity scores. The file also lists membership of 13 trios with carrier offspring assessed for family-based comparisons in this study.

**Table S2.** This file summarizes the number of genomic variants (SNVs, CNVs, and STRs) present in each individual in the 16p12.1 deletion cohort.

**Table S3.** This file lists Coriell Institute accession numbers for the LCL samples used in this study.

**Table S4.** This file lists differentially expressed transcripts between carriers and noncarriers of the 16p12.1 deletion, using both the main analysis and relatedness correction methods. It also includes enrichment of differentially expressed genes for Gene Ontology terms, candidate neurodevelopmental-associated genes, and genes preferentially expressed in GTEx and BrainSpan datasets.

**Table S5.** This file contains all information on the statistic tests performed in the manuscript, including sample sizes, test statistics, log-odds ratios, confidence intervals, p-values, and Benjamini-Hochberg corrected FDR. * indicates p<0.05 without multiple testing correction, and ** indicated FDR<0.05 after correction.

**Table S6.** This file lists significantly up- or down-regulated Gene Ontology biological process terms in 16p12.1 deletion carriers, as identified using Parametric Analysis of Gene Set Enrichment (PAGE).

**Table S7.** This file lists module assignments for genes derived from weighted gene co-expression network analysis, and the enrichment of genes in six modules that correspond to deletion carrier status for Gene Ontology terms.

**Table S8.** This file summarizes the numbers of gene expression changes, by family-specific pattern where applicable, identified in each individual in the 16p12.1 deletion cohort. Boxes shaded grey and labeled N/A indicate samples without available family-specific patterns for expression changes.

**Table S9.** This file lists differentially expressed genes identified in each of the offspring in all trios (n=13 with carrier offspring and n=4 with noncarrier offspring) by family-specific pattern (unique occurrence or shared with a parent), and the enrichment of each gene set for Gene Ontology terms.

**Table S10.** This file lists all genes in each individual that showed any gene expression change (differential expression, outlier expression, alternative splicing, ASE, or eQTL minor allele), with family-specific patterns when applicable, alongside the number of identified rare variants disrupting each gene.

**Table S11.** This file lists all outlier genes identified in each individual in the 16p12.1 deletion cohort, along with their expression z-scores, preferential expression in the human brain, and pLI and RVIS intolerance to variation scores.

**Table S12.** This file lists rare second-hit variants that may contribute to synergistic gene expression changes along with the 16p12.1 deletion in carrier children.

**Table S13.** This file lists isoforms and genes with alternative splicing identified in offspring of all trios (n=13 with carrier offspring and n=4 with noncarrier offspring) by family-specific pattern (unique occurrence or shared with a parent).

**Table S14.** This file lists genes with allele-specific expression identified in all individuals in the cohort, including the presence of rare deleterious coding variants on the overexpressed haplotype of each gene.

**Table S15.** This file lists identified eQTL variants in the 16p12.1 deletion cohort, including beta and FDR values, population frequency, associated eGene, and presence in GTEx LCL datasets. The file also lists all individuals in the cohort who carry a minor allele for the identified eQTLs.

## REFERENCES

1. Eichler EE. Genetic Variation, Comparative Genomics, and the Diagnosis of Disease. N Engl J Med. 2019 Jul 4;381(1):64–74.

2. Berkovic SF, Scheffer IE, Petrou S, Delanty N, Dixon-Salazar TJ, Dlugos DJ, et al. A roadmap for precision medicine in the epilepsies. Lancet Neurol. 2015 Dec 1;14(12):1219–28.

3. Schaaf CP, Betancur C, Yuen RKC, Parr JR, Skuse DH, Gallagher L, et al. A framework for an evidence-based gene list relevant to autism spectrum disorder. Nat Rev Genet. 2020 Jun 1;21(6):367–76.

4. Kavanagh DH, Tansey KE, O’Donovan MC, Owen MJ. Schizophrenia genetics: Emerging themes for a complex disorder. Mol Psychiatry. 2015 Feb 5;20(1):72–6.

5. Grove J, Ripke S, Als TD, Mattheisen M, Walters RK, Won H, et al. Identification of common genetic risk variants for autism spectrum disorder. Nat Genet. 2019;51(3):431– 44.

6. Niemi MEK, Martin HC, Rice DL, Gallone G, Gordon S, Kelemen M, et al. Common genetic variants contribute to risk of rare severe neurodevelopmental disorders. Nature. 2018 Oct 11;562(7726):268–71.

7. Manolio TA, Collins FS, Cox NJ, Goldstein DB, Hindorff LA, Hunter DJ, et al. Finding the missing heritability of complex diseases. Nature. 2009 Oct 8;461(7265):747–53.

8. Boyle EA, Li YI, Pritchard JK. An Expanded View of Complex Traits: From Polygenic to Omnigenic. Cell. 2017 Jun;169(7):1177–86.

9. Coe BP, Girirajan S, Eichler EE. A genetic model for neurodevelopmental disease. Curr Opin Neurobiol. 2012 Oct;22(5):829–36.

10. Girirajan S, Rosenfeld JA, Cooper GM, Antonacci F, Siswara P, Itsara A, et al. A recurrent 16p12.1 microdeletion supports a two-hit model for severe developmental delay. Nat Genet. 2010 Mar 14;42(3):203–9.

11. Girirajan S, Rosenfeld JA, Coe BP, Parikh S, Friedman N, Goldstein A, et al. Phenotypic Heterogeneity of Genomic Disorders and Rare Copy-Number Variants. N Engl J Med. 2012 Oct 4;367(14):1321–31.

12. Rees E, Kendall K, Pardiñas AF, Legge SE, Pocklington A, Escott-Price V, et al. Analysis of intellectual disability copy number variants for association with schizophrenia. JAMA Psychiatry. 2016 Sep 1;73(9):963–9.

13. Mefford HC, Muhle H, Ostertag P, von Spiczak S, Buysse K, Baker C, et al. Genome-wide copy number variation in epilepsy: novel susceptibility loci in idiopathic generalized and focal epilepsies. PLoS Genet. 2010 May 20;6(5):e1000962.

14. Stefansson H, Meyer-Lindenberg A, Steinberg S, Magnusdottir B, Morgen K, Arnarsdottir S, et al. CNVs conferring risk of autism or schizophrenia affect cognition in controls. Nature. 2014;505(7483):361–6.

15. Girirajan S, Eichler EE. Phenotypic variability and genetic susceptibility to genomic disorders. Hum Mol Genet. 2010 Oct 15;19(R2):R176–87.

16. Pizzo L, Jensen M, Polyak A, Rosenfeld JA, Mannik K, Krishnan A, et al. Rare variants in the genetic background modulate cognitive and developmental phenotypes in individuals carrying disease-associated variants. Genet Med. 2019;21(4):816–25.

17. Merla G, Howald C, Henrichsen CN, Lyle R, Wyss C, Zabot MT, et al. Submicroscopic deletion in patients with Williams-Beuren syndrome influences expression levels of the nonhemizygous flanking genes. Am J Hum Genet. 2006;79(2):332–41.

18. Luo R, Sanders SJ, Tian Y, Voineagu I, Huang N, Chu SH, et al. Genome-wide transcriptome profiling reveals the functional impact of rare de novo and recurrent CNVs in autism spectrum disorders. Am J Hum Genet. 2012 Jul 13;91(1):38–55.

19. Blumenthal I, Ragavendran A, Erdin S, Klei L, Sugathan A, Guide JR, et al. Transcriptional consequences of 16p11.2 deletion and duplication in mouse cortex and multiplex autism families. Am J Hum Genet. 2014 Jun 5;94(6):870–83.

20. Frésard L, Smail C, Ferraro NM, Teran NA, Li X, Smith KS, et al. Identification of rare-disease genes using blood transcriptome sequencing and large control cohorts. Nat Med. 2019 Jun 1;25(6):911–9.

21. Gonorazky HD, Naumenko S, Ramani AK, Nelakuditi V, Mashouri P, Wang P, et al. Expanding the Boundaries of RNA Sequencing as a Diagnostic Tool for Rare Mendelian Disease. Am J Hum Genet. 2019 Mar 7;104(3):466–83.

22. Mohammadi P, Castel SE, Cummings BB, Einson J, Sousa C, Hoffman P, et al. Genetic regulatory variation in populations informs transcriptome analysis in rare disease. Science. 2019 Oct 18;366(6463):351–6.

23. Pala M, Zappala Z, Marongiu M, Li X, Davis JR, Cusano R, et al. Population- and individual-specific regulatory variation in Sardinia. Nat Genet. 2017 May 1;49(5):700–7.

24. De Vries BBA, White SM, Knight SJL, Regan R, Homfray T, Young ID, et al. Clinical studies on submicroscopic subtelomeric rearrangements: A checklist. J Med Genet. 2001;38(3):145–50.

25. Bolger AM, Lohse M, Usadel B. Trimmomatic: A flexible trimmer for Illumina sequence data. Bioinformatics. 2014 Aug 1;30(15):2114–20.

26. Li H, Durbin R. Fast and accurate long-read alignment with Burrows-Wheeler transform. Bioinformatics. 2010 Jul 15;26(5):589–95.

27. Li H, Handsaker B, Wysoker A, Fennell T, Ruan J, Homer N, et al. The Sequence Alignment/Map format and SAMtools. Bioinformatics. 2009 Aug 15;25(16):2078–9.

28. Poplin R, Ruano-Rubio V, DePristo MA, Fennell TJ, Carneiro MO, Auwera GA Van der, et al. Scaling accurate genetic variant discovery to tens of thousands of samples. bioRxiv. 2017 Jul 24;201178.

29. Wang K, Li M, Hakonarson H. ANNOVAR: Functional annotation of genetic variants from high-throughput sequencing data. Nucleic Acids Res. 2010 Sep 1;38(16):e164.

30. Krumm N, Turner TN, Baker C, Vives L, Mohajeri K, Witherspoon K, et al. Excess of rare, inherited truncating mutations in autism. Nat Genet. 2015 Jun 11;47(6):582–8.

31. Karczewski KJ, Francioli LC, Tiao G, Cummings BB, Alföldi J, Wang Q, et al. The mutational constraint spectrum quantified from variation in 141,456 humans. Nature. 2020 May 28;581(7809):434–43.

32. Schwarz JM, Cooper DN, Schuelke M, Seelow D. MutationTaster2: Mutation prediction for the deep-sequencing age. Nat Methods. 2014;11(4):361–2.

33. Ernst J, Kheradpour P, Mikkelsen TS, Shoresh N, Ward LD, Epstein CB, et al. Mapping and analysis of chromatin state dynamics in nine human cell types. Nature. 2011 May 5;473(7345):43–9.

34. Kircher M, Witten DM, Jain P, O’roak BJ, Cooper GM, Shendure J. A general framework for estimating the relative pathogenicity of human genetic variants. Nat Genet. 2014 Mar 2;46(3):310–5.

35. Wang K, Li M, Hadley D, Liu R, Glessner J, Grant SFA, et al. PennCNV: An integrated hidden Markov model designed for high-resolution copy number variation detection in whole-genome SNP genotyping data. Genome Res. 2007 Nov;17(11):1665–74.

36. Abyzov A, Urban AE, Snyder M, Gerstein M. CNVnator: An approach to discover, genotype, and characterize typical and atypical CNVs from family and population genome sequencing. Genome Res. 2011 Jun;21(6):974–84.

37. Rausch T, Zichner T, Schlattl A, Stütz AM, Benes V, Korbel JO. DELLY: Structural variant discovery by integrated paired-end and split-read analysis. Bioinformatics. 2012;28(18):333–9.

38. Layer RM, Chiang C, Quinlan AR, Hall IM. LUMPY: A probabilistic framework for structural variant discovery. Genome Biol. 2014 Jun 26;15(6):R84.

39. Chen X, Schulz-Trieglaff O, Shaw R, Barnes B, Schlesinger F, Källberg M, et al. Manta: Rapid detection of structural variants and indels for germline and cancer sequencing applications. Bioinformatics. 2016;32(8):1220–2.

40. Alkan C, Coe BP, Eichler EE. Genome structural variation discovery and genotyping. Nat Rev Genet. 2011 May;12(5):363–76.

41. Pounraja VK, Jayakar G, Jensen M, Kelkar N, Girirajan S. A machine-learning approach for accurate detection of copy number variants from exome sequencing. Genome Res. 2019 Jul 1;29(7):1134–43.

42. Mousavi N, Shleizer-Burko S, Yanicky R, Gymrek M. Profiling the genome-wide landscape of tandem repeat expansions. Nucleic Acids Res. 2019;47(15):e90.

43. Mousavi N, Margoliash J, Pusarla N, Saini S, Yanicky R, Gymrek M. TRTools: a toolkit for genome-wide analysis of tandem repeats. Bioinformatics. 2020 Aug 17;ePub.

44. Ardlie KG, DeLuca DS, Segrè A V., Sullivan TJ, Young TR, Gelfand ET, et al. The Genotype-Tissue Expression (GTEx) pilot analysis: Multitissue gene regulation in humans. Science. 2015 May 8;348(6235):648–60.

45. Dobin A, Davis CA, Schlesinger F, Drenkow J, Zaleski C, Jha S, et al. STAR: Ultrafast universal RNA-seq aligner. Bioinformatics. 2013 Jan;29(1):15–21.

46. Wang L, Nie J, Sicotte H, Li Y, Eckel-Passow JE, Dasari S, et al. Measure transcript integrity using RNA-seq data. BMC Bioinformatics. 2016 Feb 3;17(1):58.

47. Li B, Dewey CN. RSEM: Accurate transcript quantification from RNA-Seq data with or without a reference genome. BMC Bioinformatics. 2011 Aug 4;12(1):323.

48. Deluca DS, Levin JZ, Sivachenko A, Fennell T, Nazaire MD, Williams C, et al. RNA-SeQC: RNA-seq metrics for quality control and process optimization. Bioinformatics. 2012 Jun;28(11):1530–2.

49. Gonzalez-Mantilla AJ, Moreno-De-Luca A, Ledbetter DH, Martin CL. A cross-disorder method to identify novel candidate genes for developmental brain disorders. JAMA Psychiatry. 2016 Mar 1;73(3):275–83.

50. Abrahams BS, Arking DE, Campbell DB, Mefford HC, Morrow EM, Weiss LA, et al. SFARI Gene 2.0: A community-driven knowledgebase for the autism spectrum disorders (ASDs). Mol Autism. 2013 Oct 3;4(1):36.

51. Purcell SM, Moran JL, Fromer M, Ruderfer D, Solovieff N, Roussos P, et al. A polygenic burden of rare disruptive mutations in schizophrenia. Nature. 2014 Feb 22;506(7487):185–90.

52. Firth H V., Richards SM, Bevan AP, Clayton S, Corpas M, Rajan D, et al. DECIPHER: Database of Chromosomal Imbalance and Phenotype in Humans Using Ensembl Resources. Am J Hum Genet. 2009 Apr 10;84(4):524–33.

53. Wright CF, Fitzgerald TW, Jones WD, Clayton S, McRae JF, Van Kogelenberg M, et al. Genetic diagnosis of developmental disorders in the DDD study: A scalable analysis of genome-wide research data. Lancet. 2015 Apr 4;385(9975):1305–14.

54. Robinson MD, McCarthy DJ, Smyth GK. edgeR: A Bioconductor package for differential expression analysis of digital gene expression data. Bioinformatics. 2009 Jan 1;26(1):139– 40.

55. Sun S, Zhu J, Mozaffari S, Ober C, Chen M, Zhou X. Heritability estimation and differential analysis of count data with generalized linear mixed models in genomic sequencing studies. Bioinformatics. 2019 Feb 1;35(3):487–96.

56. Purcell S, Neale B, Todd-Brown K, Thomas L, Ferreira MAR, Bender D, et al. PLINK: A tool set for whole-genome association and population-based linkage analyses. Am J Hum Genet. 2007;81(3):559–75.

57. Zhou X, Stephens M. Genome-wide efficient mixed-model analysis for association studies. Nat Genet. 2012 Jul;44(7):821–4.

58. Stegle O, Parts L, Piipari M, Winn J, Durbin R. Using probabilistic estimation of expression residuals (PEER) to obtain increased power and interpretability of gene expression analyses. Nat Protoc. 2012 Mar;7(3):500–7.

59. Young MD, Wakefield MJ, Smyth GK, Oshlack A. Gene ontology analysis for RNA-seq: accounting for selection bias. Genome Biol. 2010 Feb 4;11(2):R14.

60. The Gene Ontology Consortium. The Gene Ontology Resource: 20 years and still GOing strong. Nucleic Acids Res. 2019 Jan 8;47(D1):D330–8.

61. Miller JA, Ding S-L, Sunkin SM, Smith KA, Ng L, Szafer A, et al. Transcriptional landscape of the prenatal human brain. Nature. 2014 Apr 2;508(7495):199–206.

62. Petrovski S, Wang Q, Heinzen EL, Allen AS, Goldstein DB. Genic Intolerance to Functional Variation and the Interpretation of Personal Genomes. PLoS Genet. 2013 Aug 22;9(8):e1003709.

63. Lek M, Karczewski KJ, Minikel E V., Samocha KE, Banks E, Fennell T, et al. Analysis of protein-coding genetic variation in 60,706 humans. Nature. 2016 Aug 18;536(7616):285–91.

64. Kim S-Y, Volsky DJ. PAGE: parametric analysis of gene set enrichment. BMC Bioinformatics. 2005 Jun 8;6(1):144.

65. Langfelder P, Horvath S. WGCNA: An R package for weighted correlation network analysis. BMC Bioinformatics. 2008 Dec 29;9(1):559.

66. Soneson C, Love MI, Robinson MD. Differential analyses for RNA-seq: Transcript-level estimates improve gene-level inferences. F1000Res. 2016;4:1521.

67. Love MI, Huber W, Anders S. Moderated estimation of fold change and dispersion for RNA-seq data with DESeq2. Genome Biol. 2014 Dec 5;15(12):550.

68. Johnson WE, Li C, Rabinovic A. Adjusting batch effects in microarray expression data using empirical Bayes methods. Biostatistics. 2007 Jan;8(1):118–27.

69. Castel SE, Mohammadi P, Chung WK, Shen Y, Lappalainen T. Rare variant phasing and haplotypic expression from RNA sequencing with phASER. Nat Commun. 2016 Sep 8;7:12817.

70. Martin M, Patterson M, Garg S, O Fischer S, Pisanti N, Klau G, et al. WhatsHap: fast and accurate read-based phasing. bioRxiv. 2016 Nov 14;085050.

71. Delaneau O, Ongen H, Brown AA, Fort A, Panousis NI, Dermitzakis ET. A complete tool set for molecular QTL discovery and analysis. Nat Commun. 2017 May 18;8:15452.

72. Greene CS, Krishnan A, Wong AK, Ricciotti E, Zelaya RA, Himmelstein DS, et al. Understanding multicellular function and disease with human tissue-specific networks. Nat Genet. 2015 Jun 27;47(6):569–76.

73. Krishnan A, Zhang R, Yao V, Theesfeld CL, Wong AK, Tadych A, et al. Genome-wide prediction and functional characterization of the genetic basis of autism spectrum disorder. Nat Neurosci. 2016 Nov 1;19(11):1454–62.

74. Hagberg AA, Schult DA, Swart PJ. Exploring network structure, dynamics, and function using NetworkX. In: 7th Python in Science Conference (SciPy 2008). 2008. p. 11–5.

75. Harris CR, Millman KJ, van der Walt SJ, Gommers R, Virtanen P, Cournapeau D, et al. Array programming with NumPy. Nature. 2020 Sep 17;585(7825):357–62.

76. Virtanen P, Gommers R, Oliphant TE, Haberland M, Reddy T, Cournapeau D, et al. SciPy 1.0: fundamental algorithms for scientific computing in Python. Nat Methods. 2020 Mar 1;17(3):261–72.

77. McKinney W. Data Structures for Statistical Computing in Python. In: Proceedings of the 9th Python in Science Conference. 2010. p. 56–61.

78. Schubert J, Siekierska A, Langlois M, May P, Huneau C, Becker F, et al. Mutations in STX1B, encoding a presynaptic protein, cause fever-associated epilepsy syndromes. Nat Genet. 2014 Dec 11;46(12):1327–32.

79. Rao SSP, Huntley MH, Durand NC, Stamenova EK, Bochkov ID, Robinson JT, et al. A 3D map of the human genome at kilobase resolution reveals principles of chromatin looping. Cell. 2014 Dec 18;159(7):1665–80.

80. Werling DM, Pochareddy S, Choi J, An JY, Sheppard B, Peng M, et al. Whole-Genome and RNA Sequencing Reveal Variation and Transcriptomic Coordination in the Developing Human Prefrontal Cortex. Cell Rep. 2020 Apr 7;31(1):107489.

81. Courchesne E, Gazestani VH, Lewis NE. Prenatal Origins of ASD: The When, What, and How of ASD Development. Trends Neurosci. 2020 May 1;43(5):326–42.

82. Skene NG, Roy M, Grant SG. A genomic lifespan program that reorganises the young adult brain is targeted in schizophrenia. Elife. 2017 Sep 12;6:e17915.

83. Li X, Kim Y, Tsang EK, Davis JR, Damani FN, Chiang C, et al. The impact of rare variation on gene expression across tissues. Nature. 2017 Oct 11;550(7675):239–43.

84. Zhao J, Akinsanmi I, Arafat D, Cradick TJ, Lee CM, Banskota S, et al. A Burden of Rare Variants Associated with Extremes of Gene Expression in Human Peripheral Blood. Am J Hum Genet. 2016 Feb 4;98(2):299–309.

85. Castel SE, Cervera A, Mohammadi P, Aguet F, Reverter F, Wolman A, et al. Modified penetrance of coding variants by cis-regulatory variation contributes to disease risk. Nat Genet. 2018 Sep 1;50(9):1327–34.

86. Ballouz S, Dörfel M, Crow M, Crain J, Faivre L, Keegan CE, et al. Not by systems alone: Replicability assessment of disease expression signals. bioRxiv. 2017 Apr 18;128439.

87. Mao D, Reuter CM, Ruzhnikov MRZ, Beck AE, Farrow EG, Emrick LT, et al. De novo EIF2AK1 and EIF2AK2 Variants Are Associated with Developmental Delay, Leukoencephalopathy, and Neurologic Decompensation. Am J Hum Genet. 2020 Apr 2;106(4):570–83.

88. Yuan H, Zhang L, Chen M, Zhu J, Meng Z, Liang L. A de novo triplication on 2q22.3 including the entire ZEB2 gene associated with global developmental delay, multiple congenital anomalies and behavioral abnormalities. Mol Cytogenet. 2015 Dec 23;8(1):99.

89. Baxter AL, Vivian JL, Hagelstrom RT, Hossain W, Golden WL, Wassman ER, et al. A Novel Partial Duplication of ZEB2 and Review of ZEB2 Involvement in Mowat-Wilson Syndrome. Mol Syndromol. 2017 Jun 1;8(4):211–8.

90. Zahler AM, Rogel LE, Glover ML, Yitiz S, Ragle JM, Katzman S. SNRP-27, the C. Elegans homolog of the tri-snRNP 27K protein, has a role in 5′ splice site positioning in the spliceosome. RNA. 2018 Oct 1;24(10):1314–25.

91. Saito T, Guan F, Papolos DF, Lau S, Klein M, Fann CSJ, et al. Mutation analysis of SYNJ1: A possible candidate gene for chromosome 21q22-linked bipolar disorder. Mol Psychiatry. 2001;6(4):387–95.

92. Mitchel MW, Moreno-De-Luca D, Myers SM, Finucane B, Ledbetter DH, Martin CL. 17q12 Recurrent Deletion Syndrome. In: Adam MP, Ardinger HH, Pagon RA, Wallace SE, Bean LJ, Stephens K, et al., editors. GeneReviews®. University of Washington, Seattle; 2016.

93. Lim ET, Uddin M, De Rubeis S, Chan Y, Kamumbu AS, Zhang X, et al. Rates, distribution and implications of postzygotic mosaic mutations in autism spectrum disorder. Nat Neurosci. 2017 Sep 1;20(9):1217–24.

94. Ramírez-Castillejo C, Sánchez-Sánchez F, Andreu-Agulló C, Ferrón SR, Aroca-Aguilar JD, Sánchez P, et al. Pigment epithelium-derived factor is a niche signal for neural stem cell renewal. Nat Neurosci. 2006 Mar 19;9(3):331–9.

95. Yao I, Iida J, Nishimura W, Hata Y. Synaptic and nuclear localization of brain-enriched guanylate kinase-associated protein. J Neurosci. 2002 Jul 1;22(13):5354–64.

96. De Wit MCY, De Coo IFM, Halley DJJ, Lequin MH, Mancini GMS. Movement disorder and neuronal migration disorder due to ARFGEF2 mutation. Neurogenetics. 2009;10(4):333–6.

97. Nowak F V. Porf-2 = arhgap39 = vilse: A pivotal role in neurodevelopment, learning and memory. eNeuro. 2018 Sep 1;5(5):e0082–18.2018.

98. Dickson SP, Wang K, Krantz I, Hakonarson H, Goldstein DB. Rare Variants Create Synthetic Genome-Wide Associations. PLoS Biol. 2010 Jan 26;8(1):e1000294.

99. Zhang X, Zhang Y, Zhu X, Purmann C, Haney MS, Ward T, et al. Local and global chromatin interactions are altered by large genomic deletions associated with human brain development. Nat Commun. 2018 Dec 1;9(1):5356.

100. Powell JE, Henders AK, McRae AF, Kim J, Hemani G, Martin NG, et al. Congruence of Additive and Non-Additive Effects on Gene Expression Estimated from Pedigree and SNP Data. PLoS Genet. 2013 May 16;9(5):e1003502.

101. Lenz TL, Deutsch AJ, Han B, Hu X, Okada Y, Eyre S, et al. Widespread non-additive and interaction effects within HLA loci modulate the risk of autoimmune diseases. Nat Genet. 2015 Aug 27;47(9):1085–90.

102. Nordsletten AE, Larsson H, Crowley JJ, Almqvist C, Lichtenstein P, Mataix-Cols D. Patterns of nonrandom mating within and across 11 major psychiatric disorders. JAMA Psychiatry. 2016 Apr 1;73(4):354–61.

103. Owen MJ. Intellectual disability and major psychiatric disorders: A continuum of neurodevelopmental causality. Br J Psychiatry. 2012 Apr;200(4):268–9.

104. Connolly S, Anney R, Gallagher L, Heron EA. Evidence of Assortative Mating in Autism Spectrum Disorder. Biol Psychiatry. 2019 Aug 15;86(4):286–93.

105. Pizzo L, Lasser M, Yusuff T, Jensen M, Ingraham P, Huber E, et al. Functional assessment of the “two-hit” model for neurodevelofpmental defects in Drosophila and X. laevis. PLoS Genet. 2021 Apr 1;17(4):e1009112.

106. Wang Y, Song F, Zhang B, Zhang L, Xu J, Kuang D, et al. The 3D Genome Browser: A web-based browser for visualizing 3D genome organization and long-range chromatin interactions. Genome Biol. 2018 Oct 4;19(1):151.

