## Supplemental Figures for "Combinatorial patterns of gene expression changes contribute to variable expressivity of the developmental delay-associated 16p12.1 deletion"

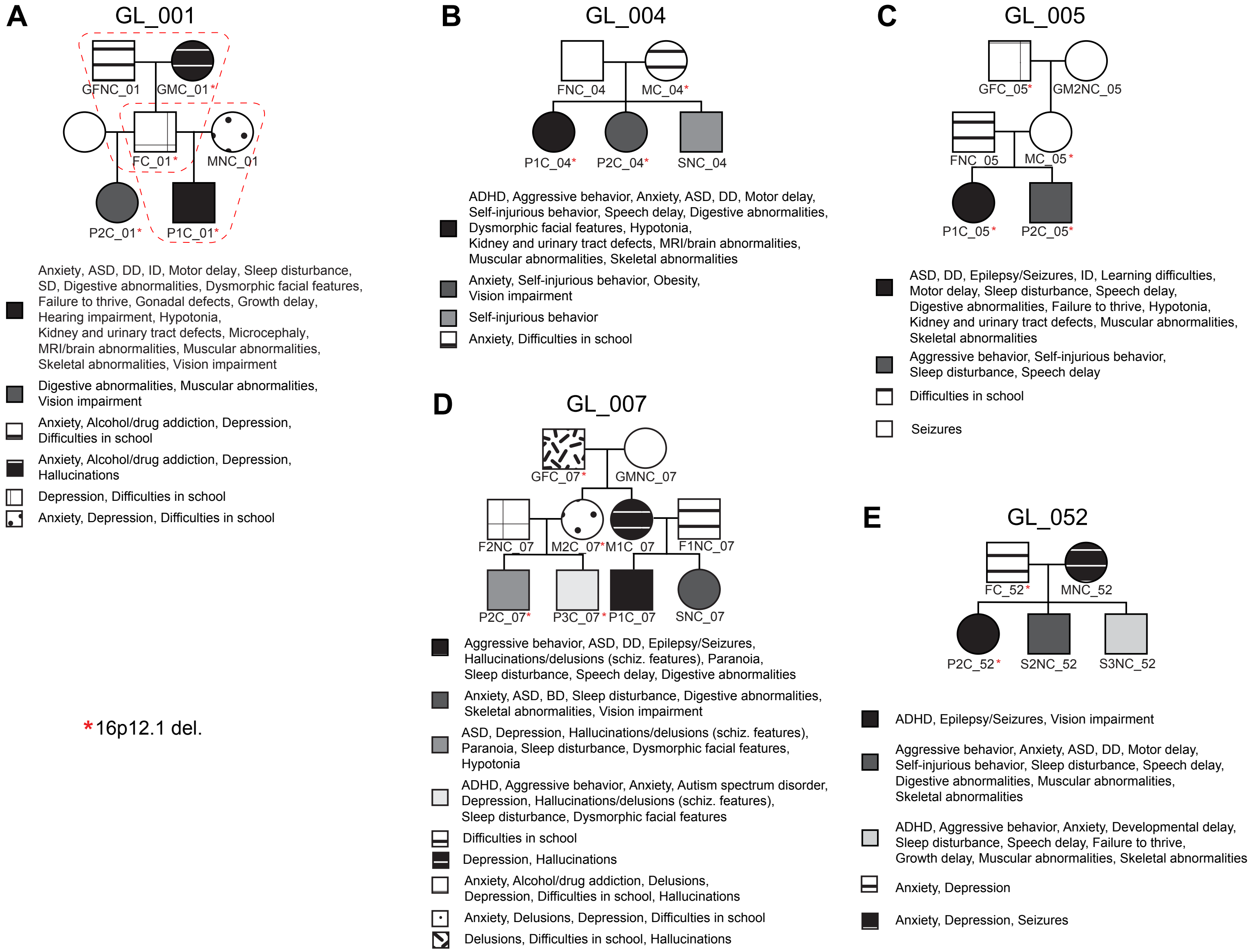


**Figure S1. Pedigrees of families with the 16p12.1 deletion.** Pedigrees of families in the 16p12.1 deletion cohort with known neuropsychiatric clinical features. * indicates carrier of 16p12.1 deletion. Dotted lines in family GL_001 indicate two trios assessed for family-based studies (**Table S1**). "ID", intellectual disability; "DD", developmental delay; "ADHD", attention deficit hyperactivity disorder; "ASD", autism spectrum disorder; "SCZ", schizophrenia; "BD", bipolar disorder.

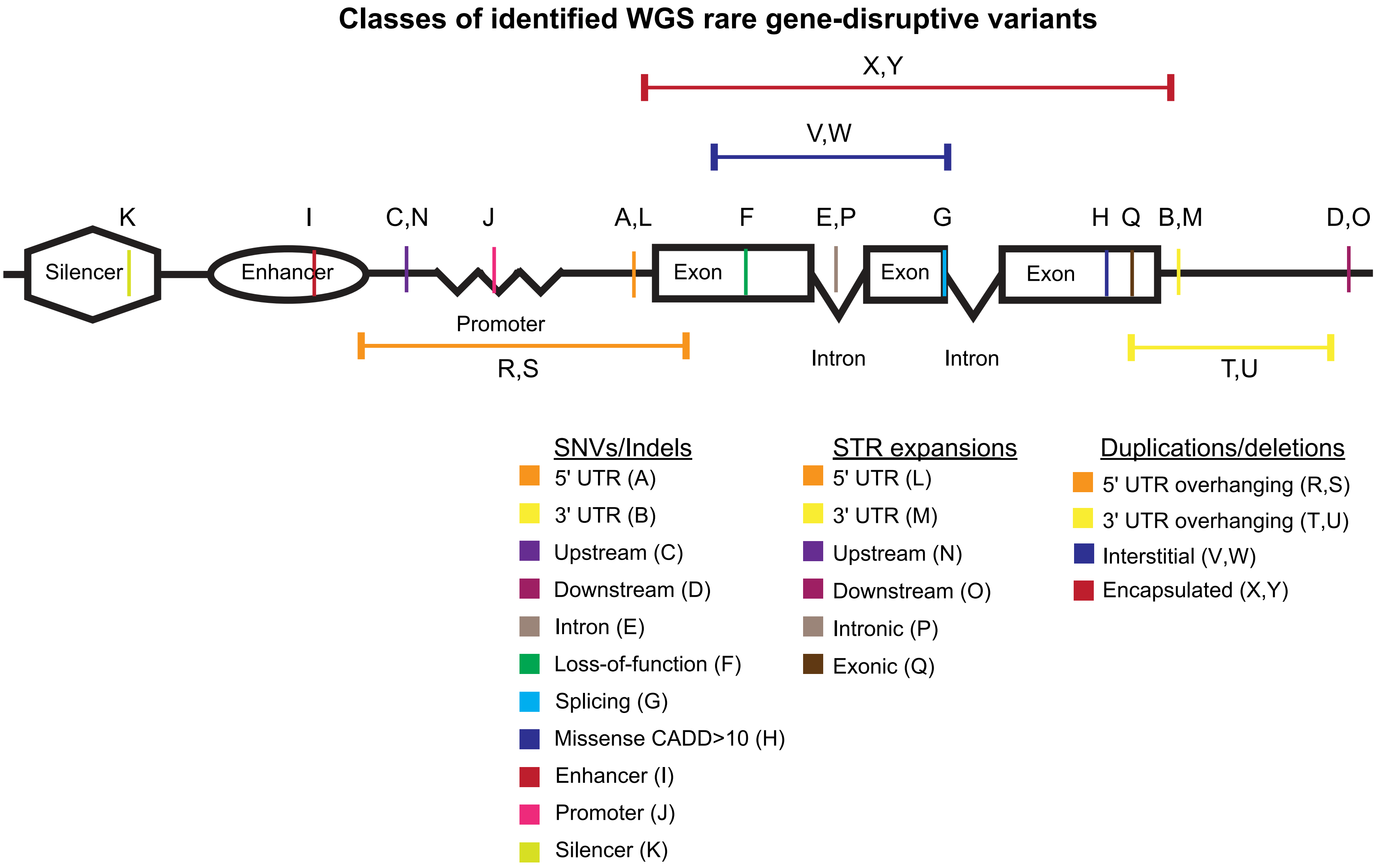


**Figure S2. Identification of rare gene-disrupting variants from WGS data.** Schematic shows the genomic locations in relation to a mock gene for 25 classes of rare coding and non-coding SNVs/indels, copy-number variants (duplications and deletions), and short tandem repeat (STR) expansions identified from sequencing data. Each variant class is color-coded and labeled by letter (A-Y). The number of rare variants in each class per individual is presented in **Table S2**.

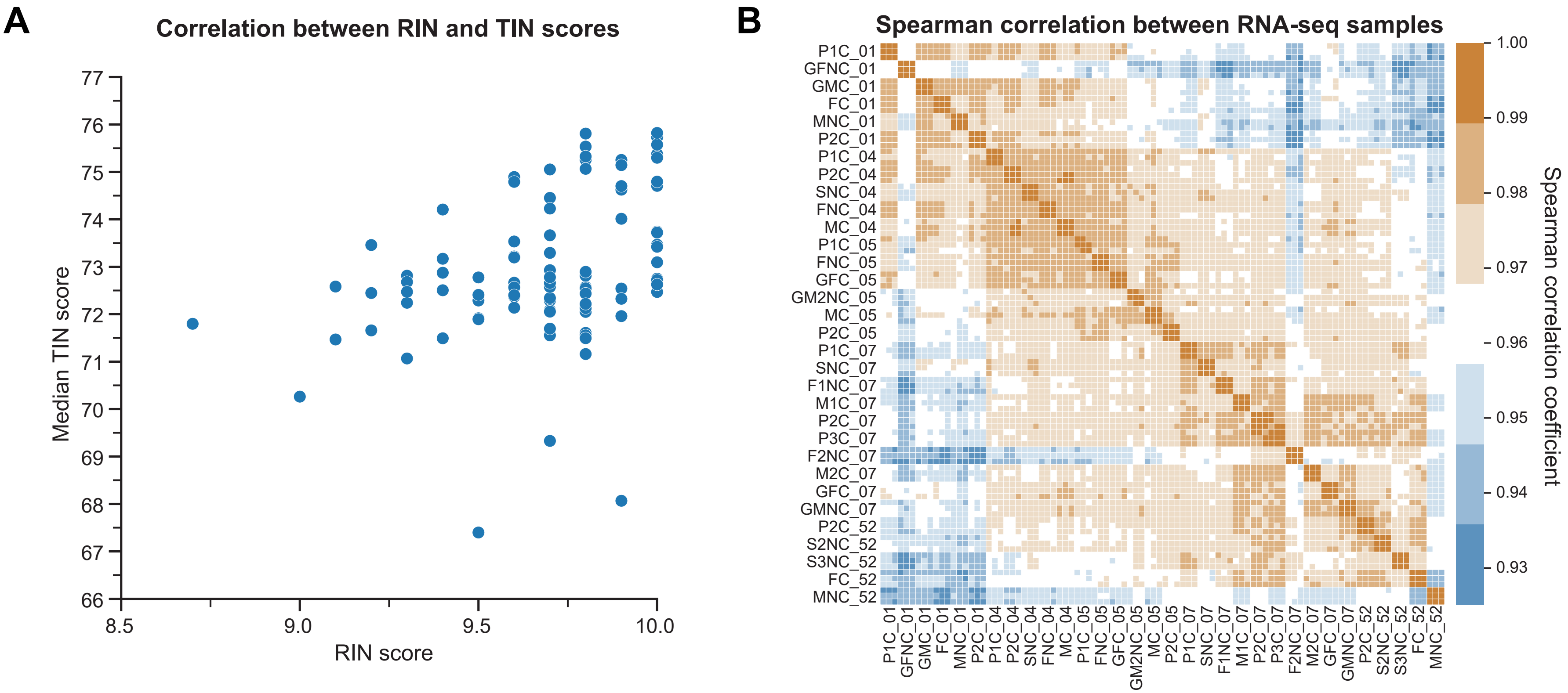


**Figure S3. Quality control of RNA sequencing data.** **(A)** Scatter plot shows moderate correlation between sample RIN scores and median TIN scores (r=0.38, p=$1.0\times10^{-4}$, Pearson correlation) of the 96 RNA-Seq replicates. **(B)** Heatmap shows Spearman correlation coefficients between pairs of 96 RNA-sequencing sample replicates. Pairs of replicates corresponding to the same individual showed higher correlation coefficients (orange) than replicates from different individuals.

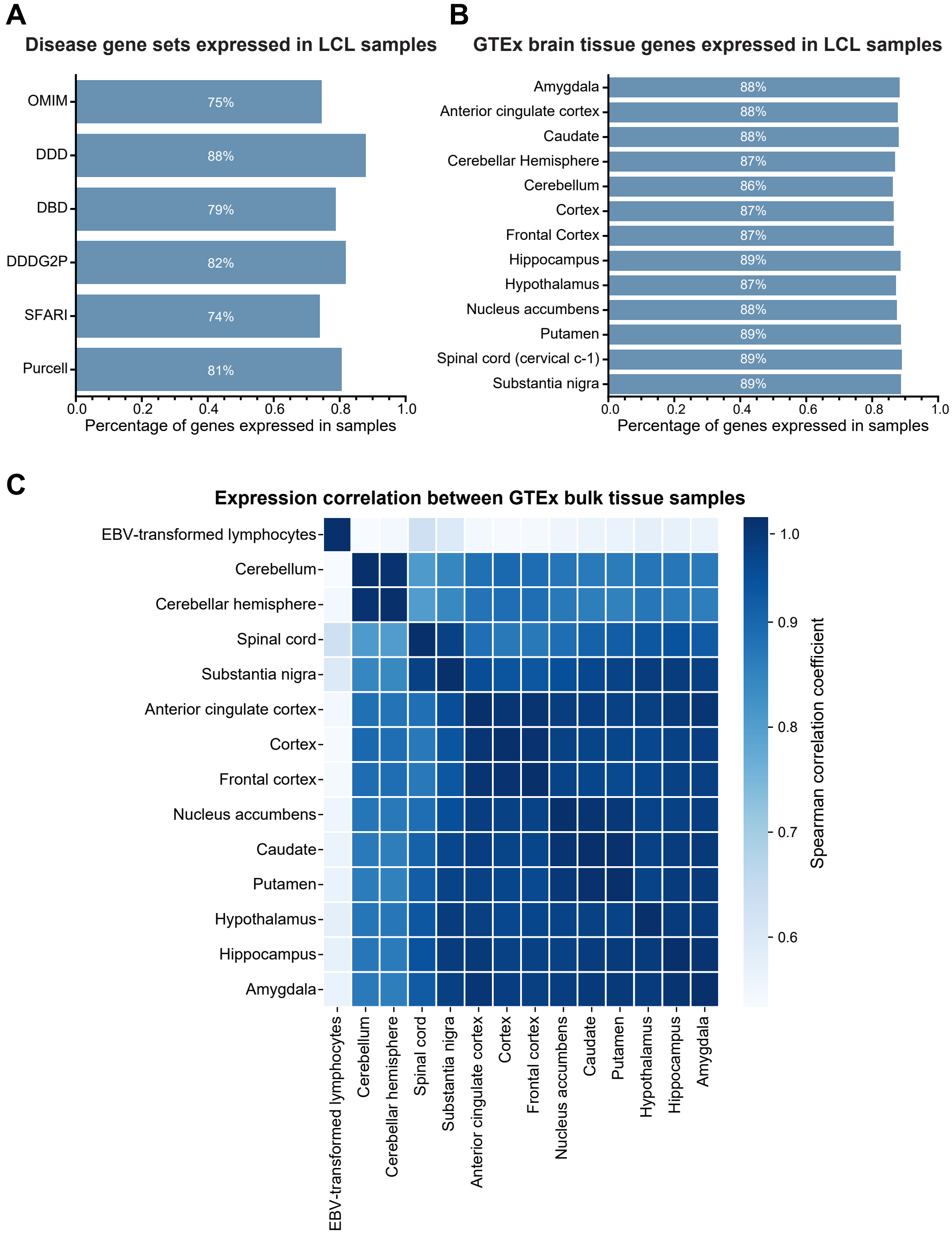


**Figure S4. Genes expressed in LCL samples by disease relevance. (A)** Percentage of genes in candidate disease gene databases that are expressed in the 32 LCL samples (>0.2 TPM in all replicates of ≥1 sample). "OMIM", Online Mendelian Inheritance in Man; "DDD", Deciphering Developmental Disorders (ID/DD); "DBD", Geisinger Developmental Brain Disorder; "DDDG2P", Developmental Disorders Genotype-Phenotype Database (ID/DD); "SFARI", Simons Foundation Autism Research Initiative; "Purcell", GeneBook from Purcell and colleagues (schizophrenia) (1). (**B)** Percentage of genes expressed in GTEx adult brain tissues that are expressed in the 32 LCL samples. GTEx expression was defined as genes that showed >0.5 TPM in 80% of samples for a particular tissue. **(C)** Heatmap depicts Spearman correlation coefficients between the median expression of adult brain tissues and lymphocyte samples from GTEx datasets (2).

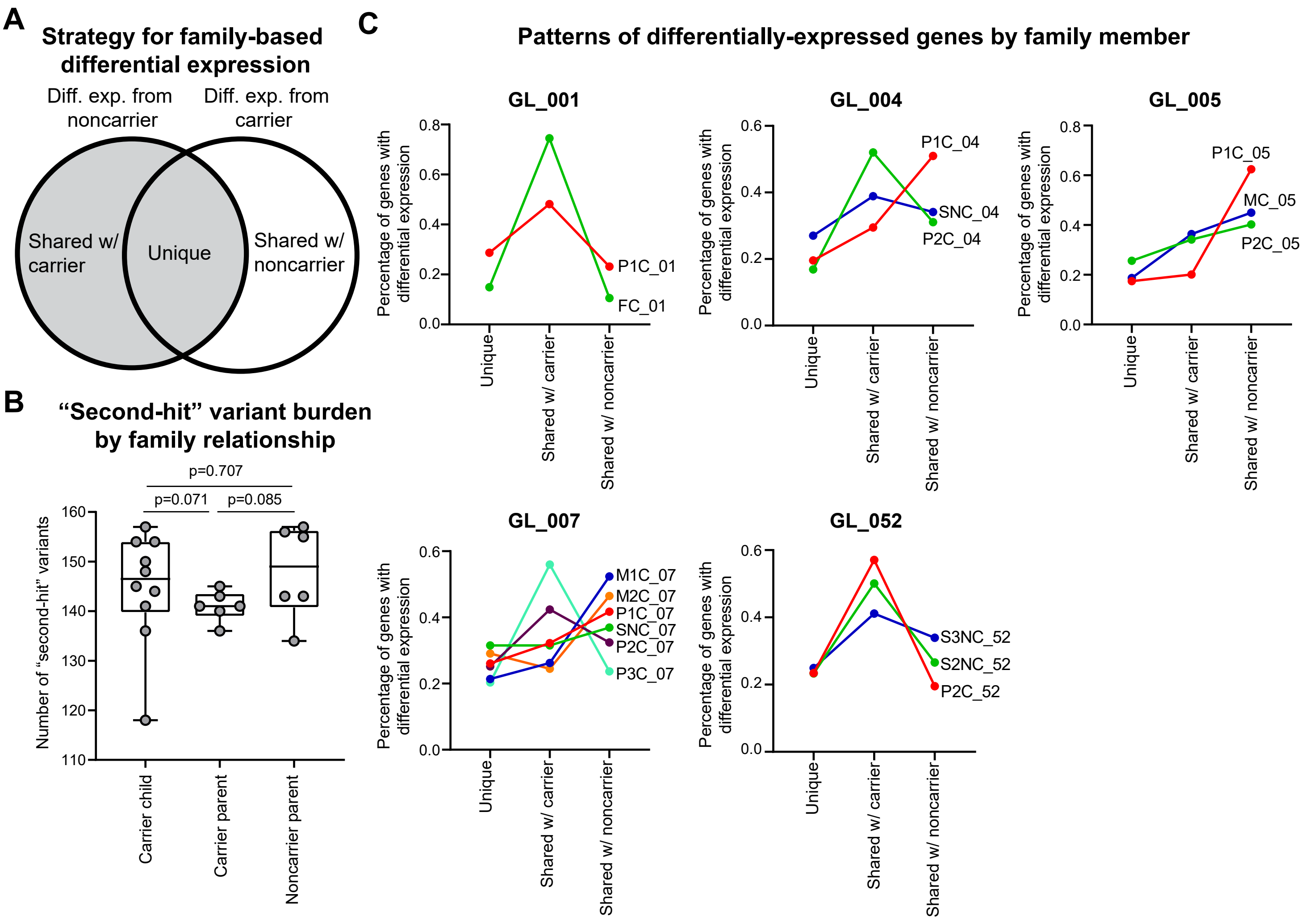


**Figure S5. Family-based differential gene expression analysis. (A)** To identify family-specific patterns for gene expression changes in 13 trios among the five families, we performed differential expression analysis between offspring and their carrier parents, and separately identified differentially expressed genes between offspring and noncarrier parents. After overlapping these two gene sets, we assigned gene expression changes in the offspring as unique (different in offspring than both parents), shared with the carrier parent (different in offspring than noncarrier parent), or shared with the noncarrier parent (different in offspring than carrier parent). **(B)** Carrier children (n=10) have a non-significant trend towards a higher burden of rare deleterious variants compared with carrier parents (n=6, p=0.071, one-tailed Mann-Whitney test), mirroring trends previously observed in a larger cohort of 26 deletion families (3). Boxplot indicates median (center line), 25th and 75th percentiles (bounds of box), and minimum and maximum (whiskers). **(C)** Line plots show family-specific patterns for differentially expressed genes in the child samples from 13 trios with carrier offspring (including carrier children compared to parents and carrier parents compared to grandparents) and four trios with noncarrier offspring in each of the five 16p12.1 deletion families. Individual samples are labeled by sample ID (**Table S1**).

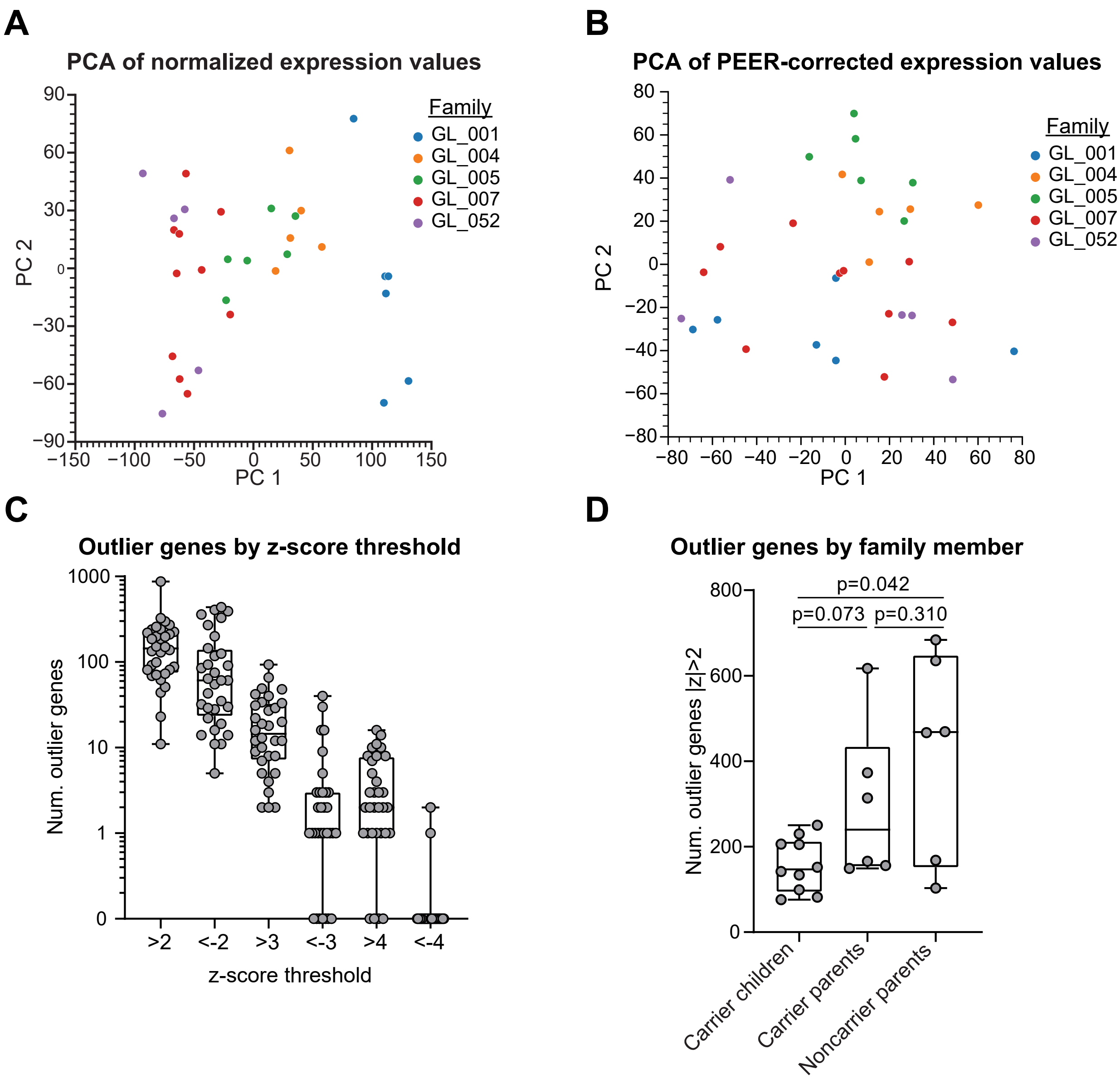


**Figure S6. Outlier gene expression analysis in the 16p12.1 deletion cohort. (A)** Scatter plot shows first two principal components (PC) of normalized expression values for RNA-seq replicates (n=96) before PEER correction. **(B)** Scatter plot shows first two PCs of PEER-corrected expression values for each RNA-Seq replicate (n=96). **(C)** Boxplot shows the number of outlier genes per individual (n=32) for different outlier expression z-score thresholds. **(D)** Boxplot shows that carrier children (n=10) have trends towards smaller numbers of genes with outlier expression (|z|>2) than carrier and noncarrier parents (n=6, p<0.073, two-tailed Mann-Whitney test). All boxplots indicate median (center line), 25th and 75th percentiles (bounds of box), and minimum and maximum (whiskers).

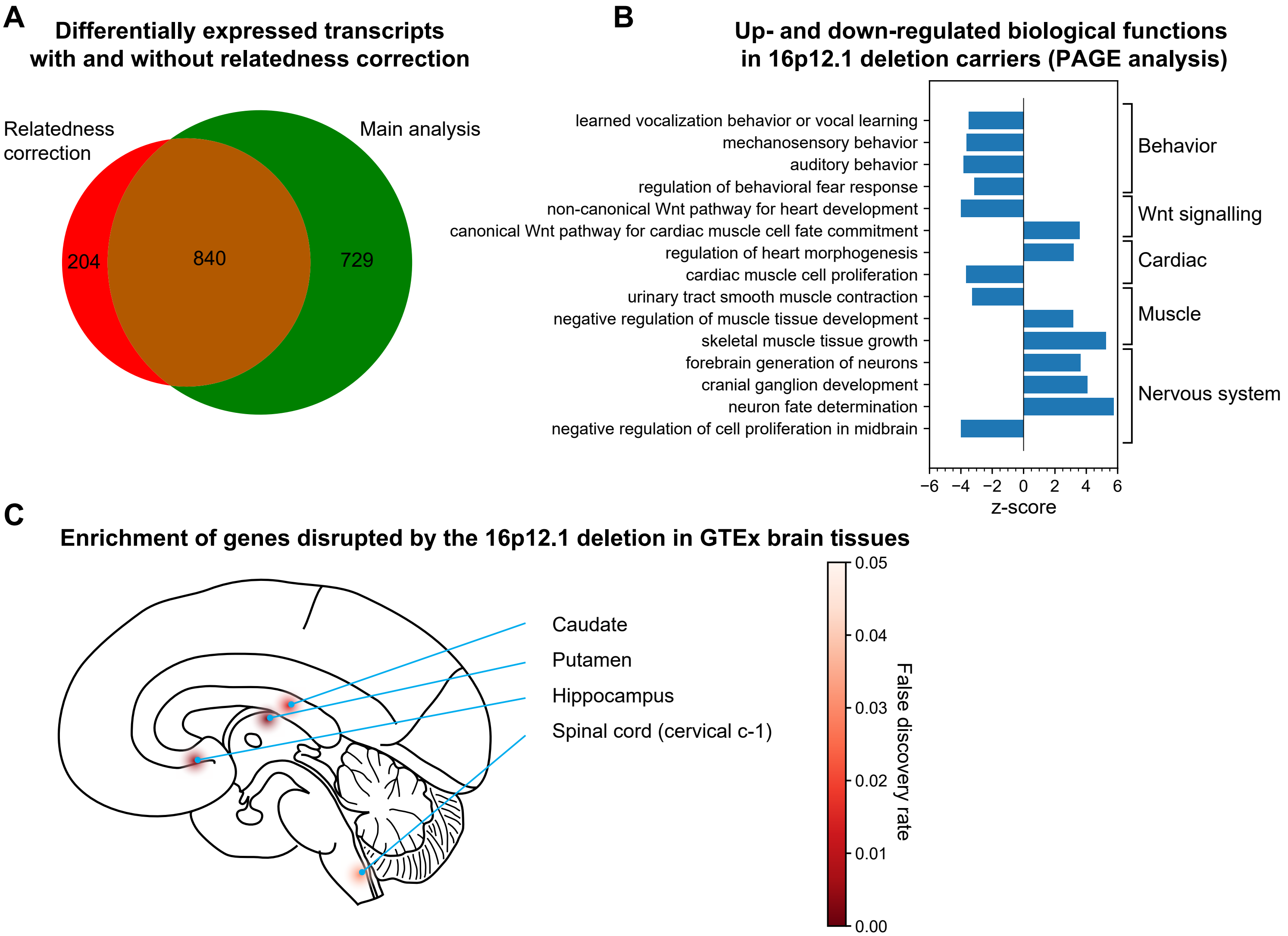


**Figure S7. Differentially expressed genes between 16p12.1 deletion carriers and noncarriers. (A)** Venn diagram shows the overlap of differentially expressed genes identified in the main analysis (using edgeR pipeline) and with relatedness correction (using PQLSeq pipeline). **(B)** Bar plot shows Gene Ontology terms whose genes are significantly up- or down-regulated in deletion carriers compared with non-carriers (PAGE analysis; FDR<0.05). Positive z-scores indicate upregulated biological functions, and negative z-scores indicate downregulated biological functions. **(C)** Diagram shows GTEx brain tissues with preferentially expressed genes that are enriched (FDR<0.05) among genes differentially expressed in 16p12.1 deletion carriers. GTEx preferential tissue expression was defined as expression >2 standard deviations than the median expression across all tissues for that gene. Brain diagram modified from WikiCommons open-source image (https://commons.wikimedia.org/wiki/File:Gehirn,_medial_-_Lobi_en.svg).

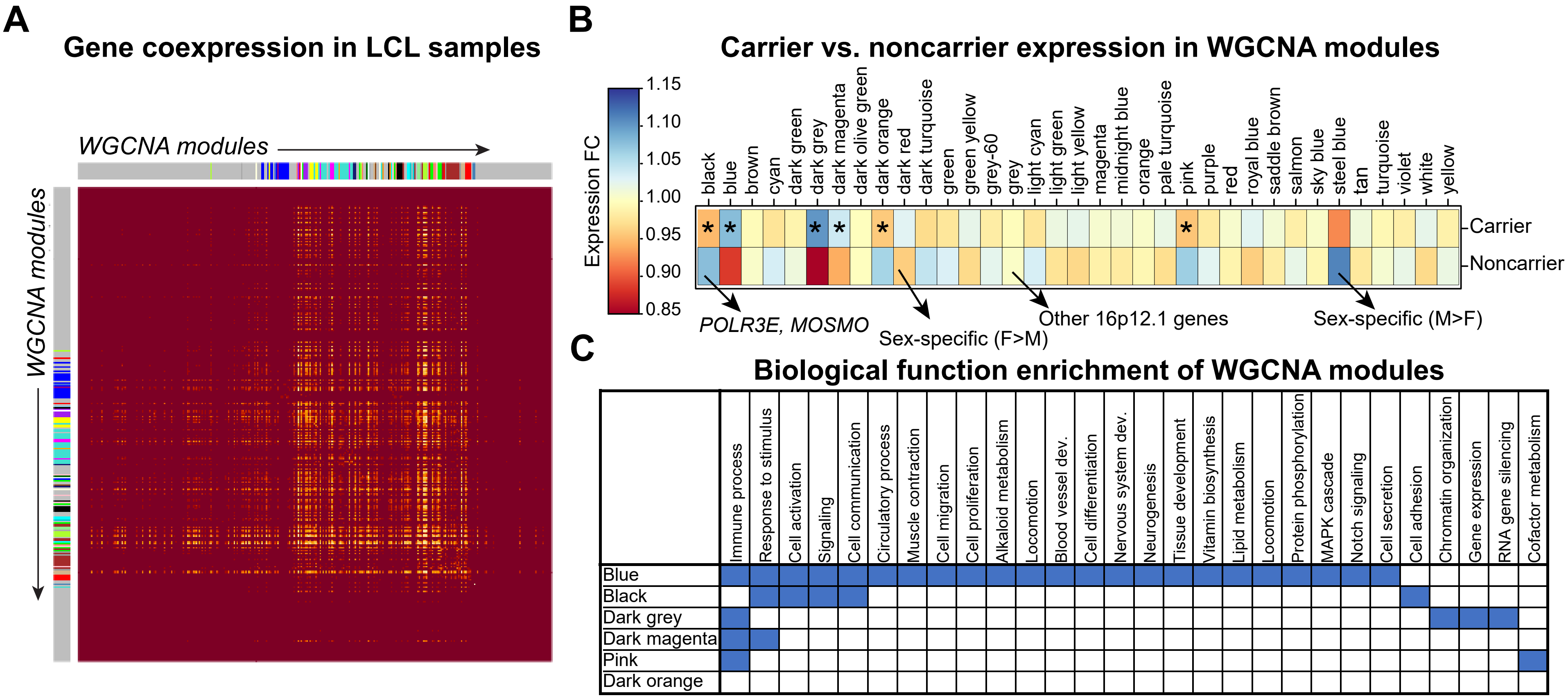


**Figure S8. Weighted gene co-expression network analysis. (A)** Heatmap showing co-expression between pairs of genes expressed in LCL samples derived from all individuals within the five 16p12.1 deletion families (n=32 samples). Yellow cells indicate pairs of genes that are co-expressed across samples, and colored bars on the side of the heatmap represent assignment of genes to one of 35 WGCNA modules. **(B)** Heatmap showing expression fold-change differences of genes in the 35 WGCNA modules between LCL samples of carriers (n=57 replicates of 19 samples) and noncarriers (n=39 replicates of 13 samples). Two modules showed sex-specific differences and were excluded for further analysis. Six modules showed significant expression differences between carriers and noncarriers (*FDR<0.05, two-tailed T-test with Benjamini-Hochberg correction). **(C)** Table showing clusters of enriched (FDR<0.05) Gene Ontology (GO) Biological Process terms in the six WGCNA modules with significant carrier-noncarrier differences. The dark orange module did not have any significant enrichments.

### Functional enrichments of differentially-expressed genes in carrier children

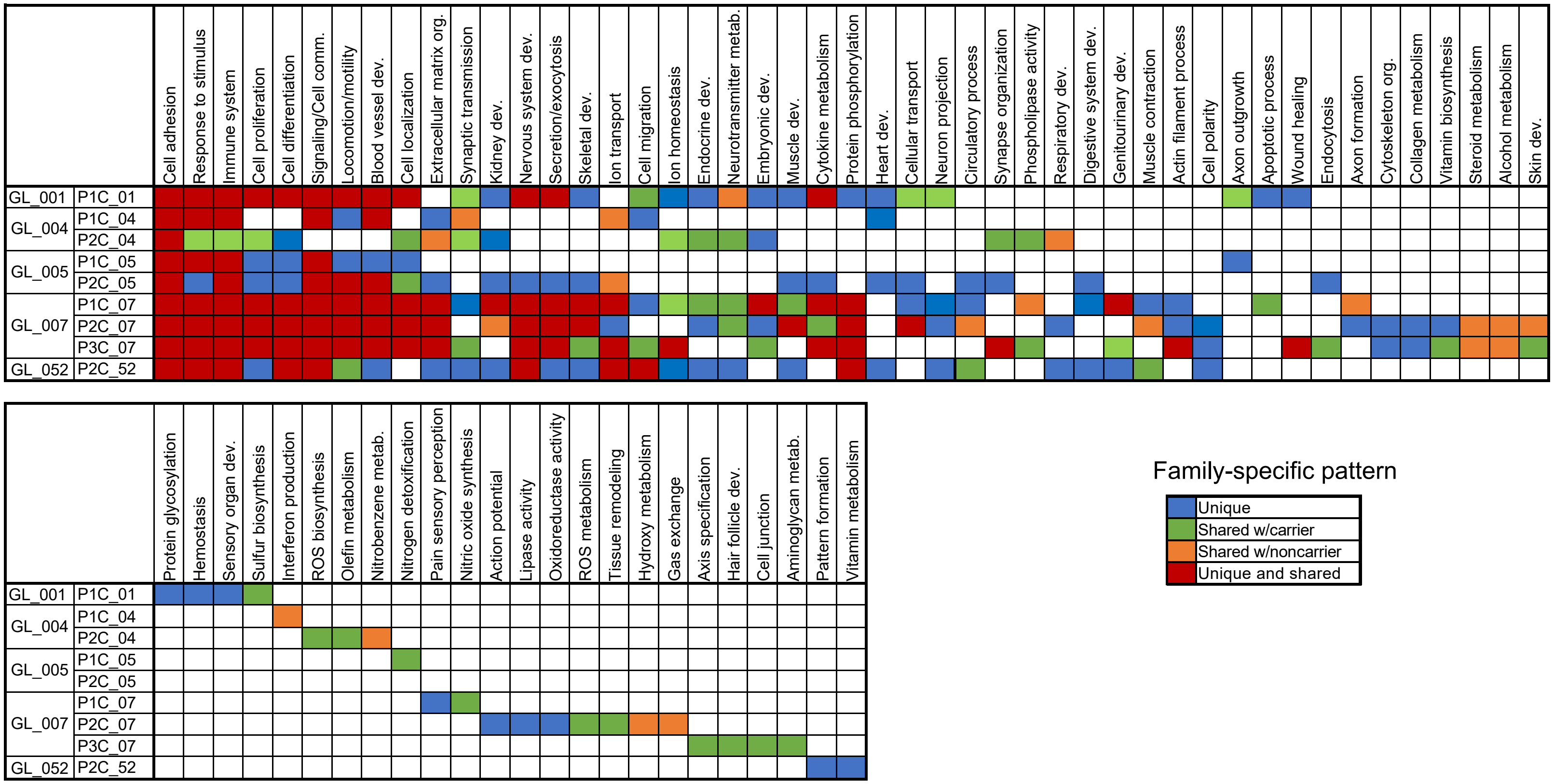


**Figure S9. Functional enrichments of differentially expressed genes in carrier children.** Table shows clusters of Gene Ontology (GO) biological function terms enriched (FDR<0.05) among differentially expressed genes in carrier children with the 16p12.1 deletion (n=9), colored by family-specific patterns for the differentially expressed gene sets (unique, shared with the carrier or noncarrier parent, or both unique and shared with either parent).

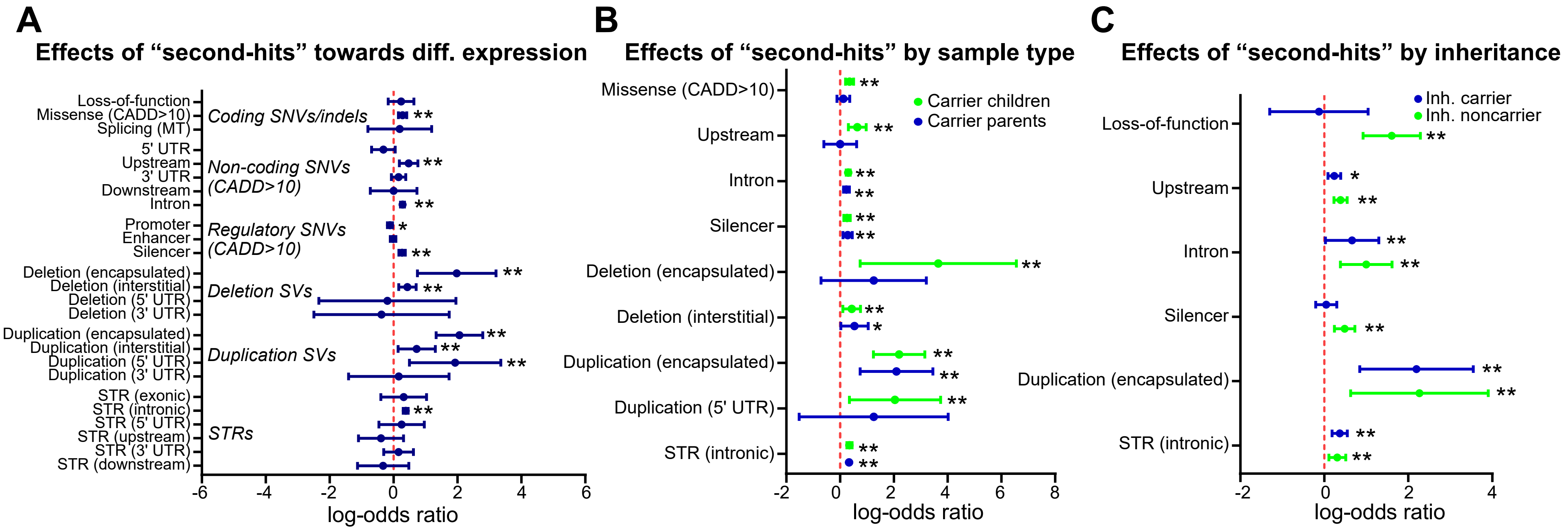


**Figure S10. Enrichment of differentially expressed genes for rare variants. (A)** Forest plot shows enrichment (Fisher's exact test, **=FDR<0.05, *=uncorrected p<0.05) of differentially expressed genes in the carrier offspring of 13 trios for proximal coding and non-coding rare variants, including single-nucleotide variants (SNVs) and insertions/deletions (indels) with CADD scores >10 (4), structural variants (SVs), and short tandem repeats (STRs). **(B)** Forest plot shows classes of rare variants with significant enrichment (Fisher's exact test, **=FDR<0.05, *=uncorrected p<0.05) towards differentially expressed genes in carrier children (n=9) or carrier parents (n=4). **(C)** Forest plot shows classes of rare variants with significant enrichment (Fisher's exact test, **=FDR<0.05, *=uncorrected p<0.05) towards differentially expressed genes in carrier children (n=9) that are shared with either carrier or noncarrier parents. All forest plots show log-odds ratios (dots) and 95% confidence intervals (whiskers). Odds ratios, confidence intervals, p-values, and Benjamini-Hochberg corrected FDRs for comparisons with all classes of “second-hit” variants are listed in **Table S5**.

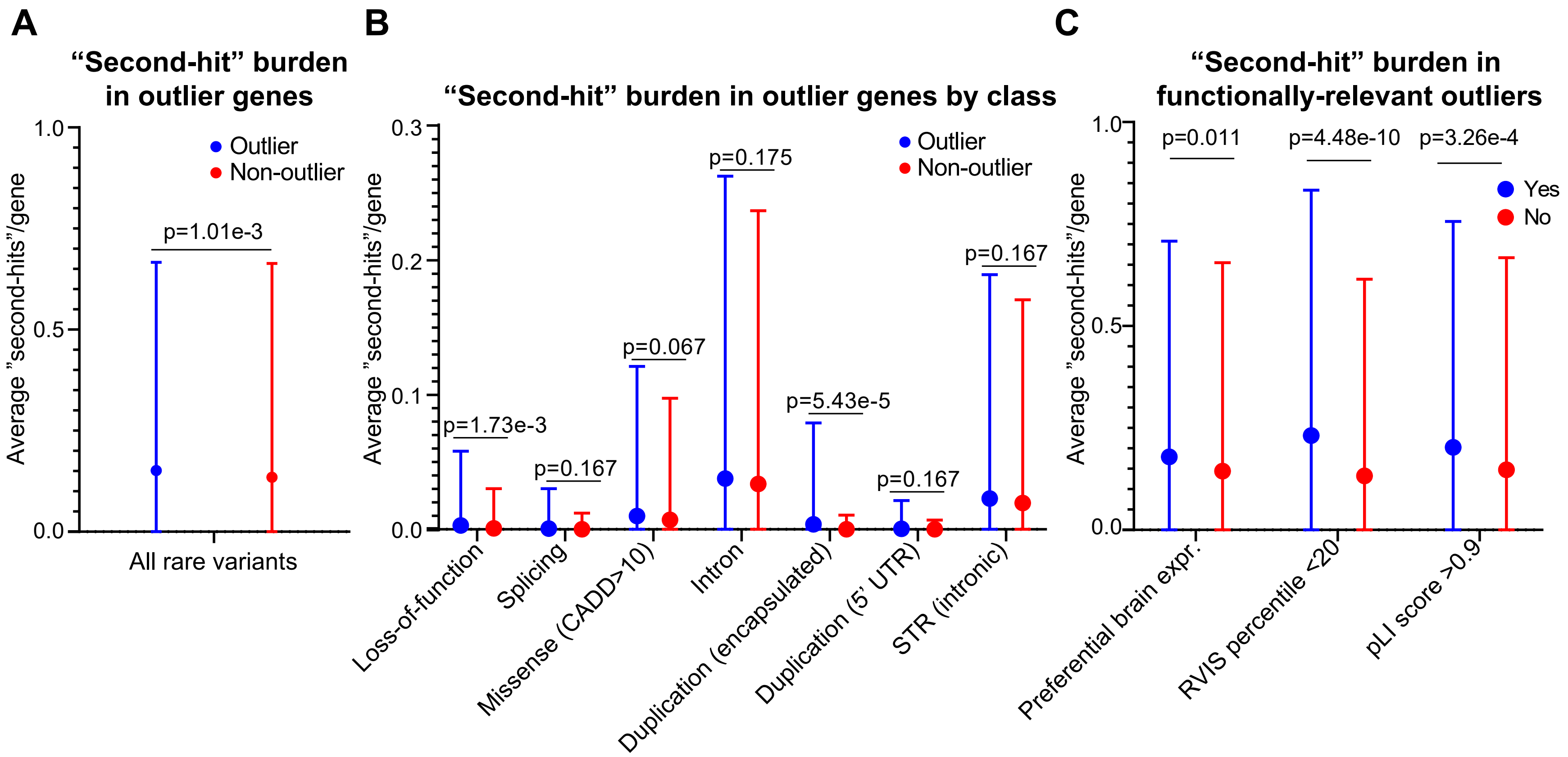


**Figure S11. Rare variant burden of outlier genes in the 16p12.1 deletion cohort. (A)** Outlier expression genes in all individuals (n=32) have a higher average burden of rare variants than non-outlier genes ($p=1.01\times10^{-3}$, one-tailed t-test). **(B)** Plots show individual classes of rare variants with increased average burden (FDR, one-tailed t-test with Benjamini-Hochberg correction) in outlier genes across all samples (n=32) compared with non-outlier genes. **(C)** Outlier genes with preferential expression in GTEx brain tissues (2), RVIS scores <20$^{th}$ percentile (5), or pLI scores >0.9 (6) have a higher average burden of rare variants than outlier genes not in those categories ($p<0.05$, two-tailed t-test) across all samples (n=32). All plots show mean +/- SD of the average burden of rare variants in each gene class.

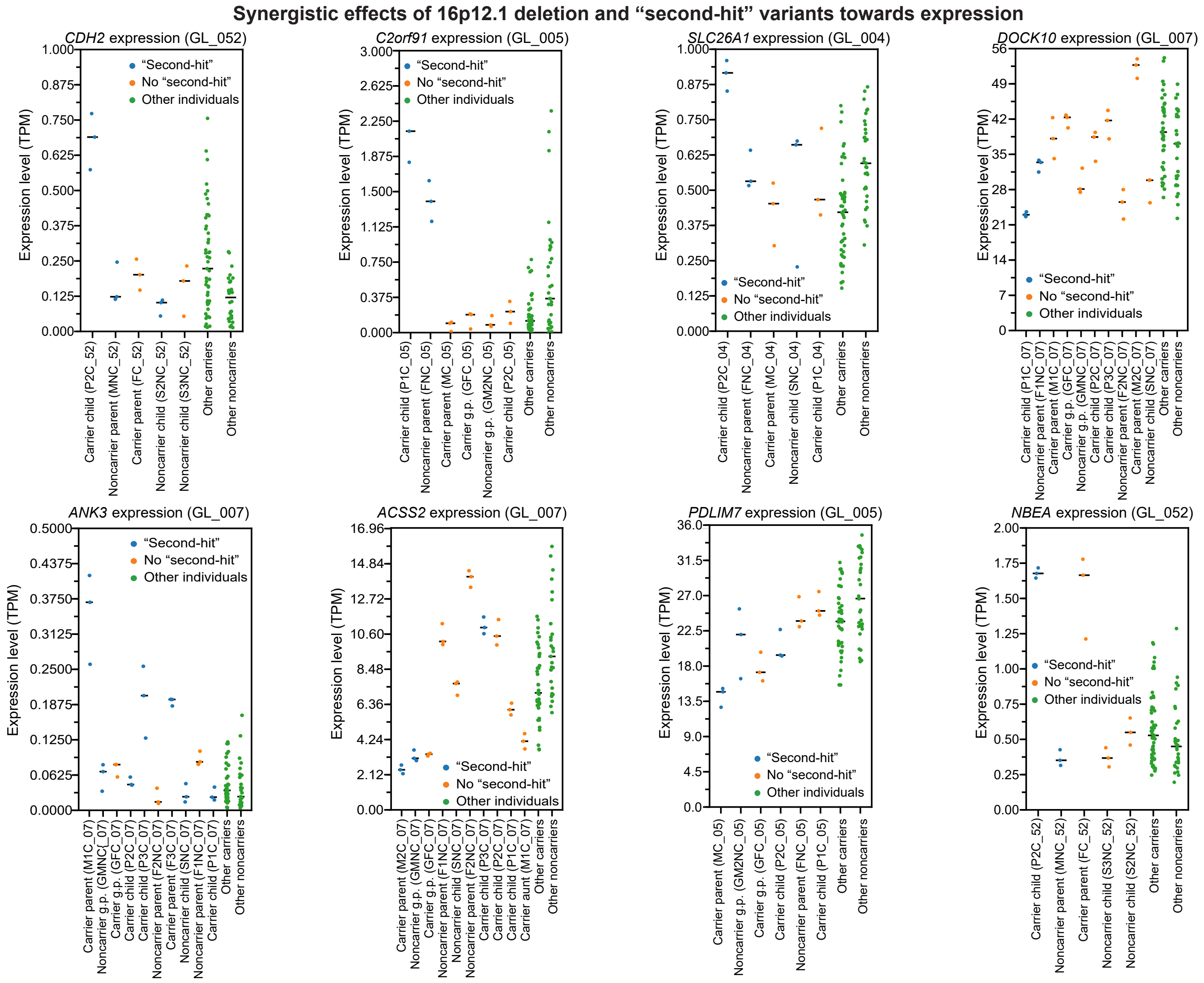


**Figure S12. Examples of synergistic effects between 16p12.1 deletion and rare variants on expression.** Scatter plots show normalized expression values (TPM) of genes with synergistic effects due to the 16p12.1 deletion and inherited “second-hit” variants (see Methods). Blue circles indicate expression values for samples from deletion carriers and family members with rare “second-hit” variants, orange circles indicate expression values for samples from family members without the rare variant, and green circles indicate expression values of samples from other deletion carriers and noncarriers in the cohort. Black lines denote median gene expression for LCL replicates of each individual used to identify genes with outlier expression in individual deletion carriers.

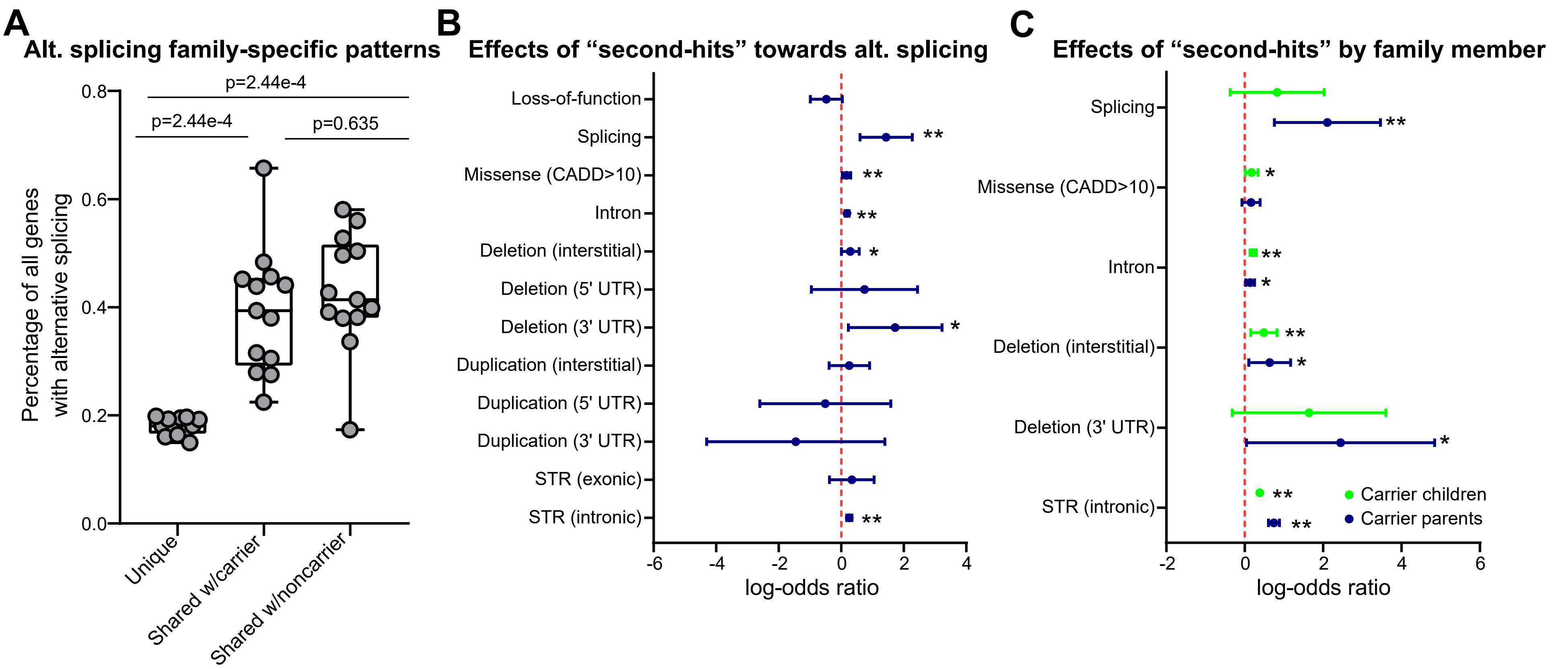


**Figure S13. Family-based alternative splicing analysis. (A)** Boxplot shows the distribution of alternative splicing events in the carrier offspring of 13 trios within 16p12.1 deletion families based on family-specific patterns. Unique events in offspring occurred at a lower frequency ($p=2.44\times10^{-4}$, two-tailed paired Mann-Whitney test) than those shared with either parent. Boxplot indicates median (center line), 25th and 75th percentiles (bounds of box), and minimum and maximum (whiskers). **(B)** Forest plot shows enrichment (Fisher's exact test, **=FDR<0.05, *=uncorrected p<0.05) of alternate splicing events in the carrier offspring of 13 trios for rare variant classes that could putatively affect gene splicing. **(C)** Forest plot shows classes of rare variants with significant enrichment (Fisher's exact test, **=FDR<0.05, *=uncorrected p<0.05) towards genes with alternative splicing in carrier children (n=9) or carrier parents (n=4). Forest plots show log-odds ratios (dots) and 95% confidence intervals (whiskers). Odds ratios, confidence intervals, p-values, and Benjamini-Hochberg corrected FDRs for comparisons with all classes of "second-hit" variants are listed in **Table S5**.

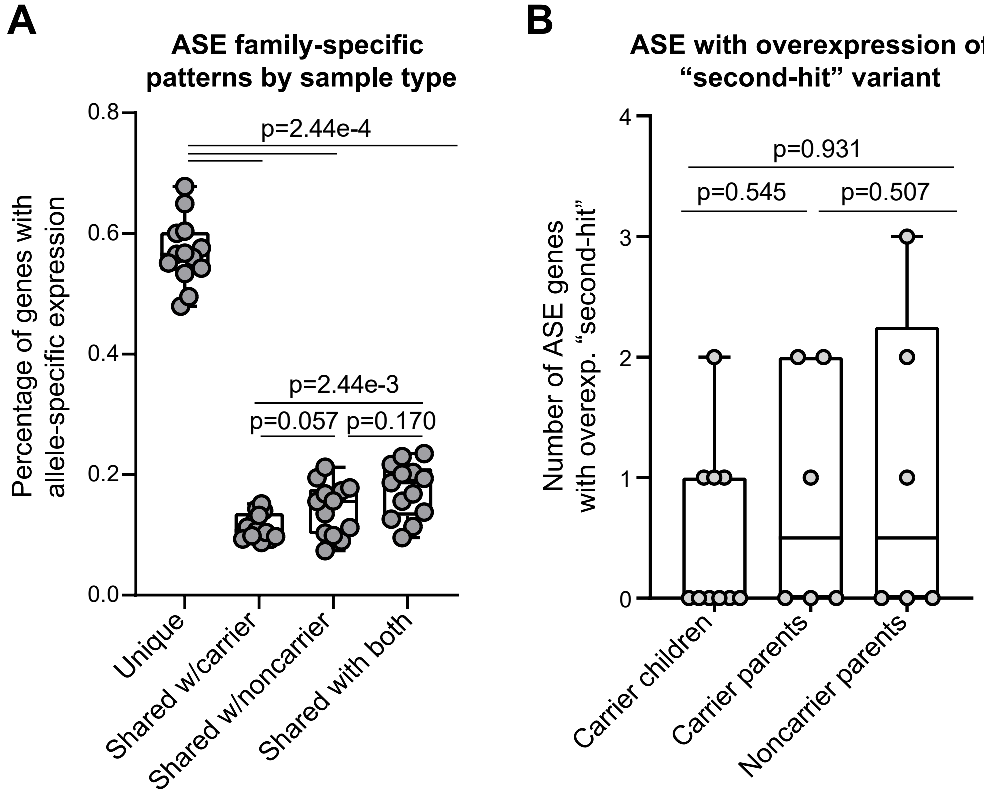


**Figure S14. Allele-specific expression analysis. (A)** Boxplot shows the distribution of allele-specific expression (ASE) events by family-specific pattern in the carrier offspring of 13 trios within the deletion families. Unique events in the offspring occurred at a higher frequency ($p=2.44\times10^{-4}$, two-tailed paired Mann-Whitney test) than events shared with either or both parents. **(B)** Boxplot shows the number of ASE events that lead to overexpression of a deleterious (CADD score>25) coding variant per individual. There were no significant differences ($p>0.05$, two-tailed Mann-Whitney test) between carrier children (n=10) and their carrier (n=6) or noncarrier parents (n=6). All boxplots indicate median (center line), 25th and 75th percentiles (bounds of box), and minimum and maximum (whiskers).

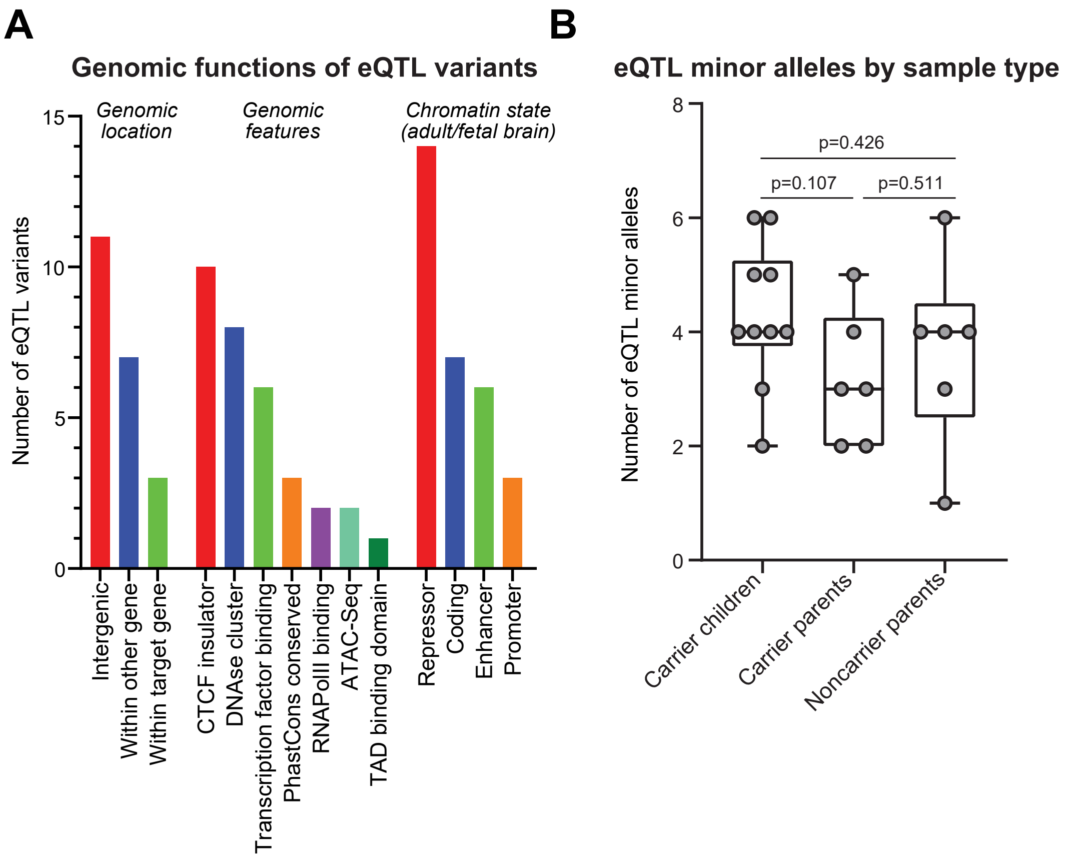


**Figure S15. Expression quantitative trait loci (eQTL) analysis. (A)** Bar plot shows the genomic characteristics of the 21 discovered eQTL variants, including genomic location, location within non-coding regulatory features, and chromatin state annotations in the adult or fetal brain. **(B)** Boxplot shows distribution of eQTL minor alleles observed in carrier children (n=10), carrier parents (n=6), and noncarrier parents (n=6). A non-significant trend (p=0.107, two-tailed Mann-Whitney test) towards higher minor allele burden was observed in carrier children compared with carrier parents. Boxplot indicates median (center line), 25th and 75th percentiles (bounds of box), and minimum and maximum (whiskers).

### SUPPLEMENTAL REFERENCES


1. Purcell SM, Moran JL, Fromer M, Ruderfer D, Solovieff N, Roussos P, et al. A polygenic burden of rare disruptive mutations in schizophrenia. Nature. 2014 Feb 22;506(7487):185–90.

2. Ardlie KG, DeLuca DS, Segrè A V., Sullivan TJ, Young TR, Gelfand ET, et al. The Genotype-Tissue Expression (GTEx) pilot analysis: Multitissue gene regulation in humans. Science. 2015 May 8;348(6235):648–60.

3. Pizzo L, Jensen M, Polyak A, Rosenfeld JA, Mannik K, Krishnan A, et al. Rare variants in the genetic background modulate cognitive and developmental phenotypes in individuals carrying disease-associated variants. Genet Med. 2019;21(4):816–25.

4. Kircher M, Witten DM, Jain P, O'roak BJ, Cooper GM, Shendure J. A general framework for estimating the relative pathogenicity of human genetic variants. Nat Genet. 2014 Mar 2;46(3):310–5.

5. Petrovski S, Wang Q, Heinzen EL, Allen AS, Goldstein DB. Genic Intolerance to Functional Variation and the Interpretation of Personal Genomes. PLoS Genet. 2013 Aug 22;9(8):e1003709.

6. Lek M, Karczewski KJ, Minikel E V., Samocha KE, Banks E, Fennell T, et al. Analysis of protein-coding genetic variation in 60,706 humans. Nature. 2016 Aug 18;536(7616):285–91.
